# PTHrP drives aggressive traits in colorectal cancer cells: Implications of tumor-stromal cells

**DOI:** 10.64898/2026.04.16.718950

**Authors:** María Belén Novoa Díaz, Pedro Carriere, Cintia Birkenstok, Stephani Gonzalez Osorio, Ariel Zwenger, Héctor R. Contreras, Claudia Gentili

## Abstract

In the tumor microenvironment (TME), dynamic interactions between cells and soluble factors promote tumor progression. We previously demonstrated that parathyroid hormone-related peptide (PTHrP), a TME-associated cytokine, enhances the aggressive phenotype of HCT116 colorectal cancer (CRC) cells, and that conditioned medium from PTHrP-treated HMEC-1 endothelial stromal cells (CM) induces epithelial-to-mesenchymal transition (EMT) in CRC cells. Here, Western blot analysis showed that CM modulates Met receptor expression and activation and promotes cancer stem cell (CSC) traits in HCT116 cells. Since PTHrP induces CPT-11 chemoresistance through Met signaling, we investigated the involvement of the CM–Met axis in this process. Viability assays revealed that CM increases cell number and confers CPT11 resistance through Met activation. Transforming growth factor beta 1 (TGFβ1), upregulated in PTHrP-treated HMEC-1 cells, was evaluated as a potential mediator. Its neutralization reversed the CM-induced increase in cell number but did not affect chemoresistance. *In silico* analyses revealed differences between CRC and normal tissues related to TGFβ1 signaling and Met activation, along with positive correlations among the analyzed markers. Immunohistochemical observation of human samples is consistent with our previous findings. Overall, these findings support a role for PTHrP in promoting CRC aggressiveness through coordinated effects on tumor and stromal compartments

## 1. Introduction

Colorectal cancer (CRC) remains a major global health burden on the global stage, as evidenced by the World Cancer Research Fund International, which indicates that CRC incidence is increasing and its prevalence is expected to rise by 60% in the next 15 years [1]. According to global cancer statistics, CRC ranks second in mortality due to emerging subtypes and resistance to therapies [1-4]. In recent decades, advances in genetic and molecular biology techniques have improved understanding of the molecular basis of CRC pathogenesis [5-7]. Guinney and collaborators proposed a categorization based on four molecular subtypes (CMS), considering key factors that contribute to the disease with special attention to the role of the tumor microenvironment (TME). The CMS1 group, also called MSI-Immune, is characterized by high microsatellite instability and immune response. CMS2 (canonical subtype) exhibits aberrant activation of the Wnt/β-catenin pathway. CMS3 (metabolic subtype) is characterized by marked deregulation of metabolic pathways, and finally, the CMS4 group (mesenchymal subtype) represents more than 20% of CRC cases and is distinguished by hyperactivation of the TGF-β signaling and marked epithelial to mesenchymal transition (EMT) [8,9]. The TME is a complex environment within several types of cells and molecules that strongly interact to create conditions for tumor growth and spread [10]. The parathyroid hormone-related peptide (PTHrP) is a cytokine present in the TME [4,11,12]. We previously observed on intestinal tumor cells that this factor induces events associated with an aggressive CRC phenotype, such as proliferation, invasion, migration, molecular and phenotypic changes related to EMT and chemoresistance [4,13,14]. Furthermore, we demonstrated that the interaction between CRC-derived cells exposed to PTHrP and HMEC-1 endothelial stromal cells, increases endothelial migratory capacity and the formation of tube-like structures [15]. It was also found that the conditioned medium derived from HMEC-1 exposed to PTHrP (TCM) favors migration and promotes the EMT program in CRC cells [13].

It is known that in the tumor context, multiple paracrine signals orchestrate key pathways that drive the synthesis and secretion of cytokines, growth factors, and other biomolecules in the TME [16,17]. Some of the most studied growth factors in tumor stroma to date are members of the TGF-β family [18]. These factors play multiple roles in cell proliferation, differentiation, apoptosis, migration, and angiogenesis and participate in different stages of neoplastic development in many types of cancer, including CRC [17,19]. Several investigations have demonstrated the interrelationship between TGF-β and PTHrP in cancer [16,17,20,21]. In osteolytic metastasis of breast cancer, the action of PTHrP releases TGF-β from the bone matrix, which stimulates the secretion of PTHrP by tumor cells, generating a cyclic mechanism [16,22]. In hepatocellular cancer, it was observed that TGF-β induces the expression and secretion of PTHrP by tumor cells, which participates in cell growth [23]. Similar behavior has been observed in prostate cancer [24] and pancreatic cancer [21]. In particular, in tissue derived from hepatocellular carcinoma, PTHrP expression was found to be predominantly present in tumor cells, while TGF-β expression was found in liver tissue adjacent to the tumor [17]. Although increasing evidence supports a functional interplay between PTHrP and TGF-β, the molecular mechanisms governing their reciprocal regulation remain poorly defined. Moreover, no studies to date have investigated this interaction in the context of CRC.

Based on the previous results and to further investigate the effect of TME on intestinal tumor cells and their interaction, we examined whether PTHrP affects crucial events in the malignant progression of CRC by regulating factors secreted by TME cells known to be involved in this disease.

## 2. Results

### 2.1. The conditioned media from HMEC-1 cells exposed to PTHrP modulates expression and activation of Met in HCT116 cells

Met is a receptor tyrosine kinase (RTK) that is frequently overexpressed and/or hyperactivated in CRC and has been implicated in tumor development and progression [25-27]. Aberrant activation of Met signaling occurs through multiple mechanisms, including receptor overexpression and/or overstimulation by its canonical ligand, hepatocyte growth factor (HGF) [28,29]. In addition, Met can be transactivated by G protein–coupled receptors (GPCRs) [30,31]. In our laboratory, we observed that CRC-derived cells express parathyroid hormone receptor type 1 (PTHR1), a GPCR [19]. The binding of PTHrP to its receptor increases Met protein expression and promotes activation of its downstream signaling pathway in our experimental model [32]. Furthermore, following xenotransplantation of intestinal tumor cells into nude mice, we observed that intratumoral administration of the cytokine increased the expression of Met and other markers associated with CRC progression [13,19,32]. Considering the direct effect of PTHrP on CRC-derived cells together with previous evidence demonstrating its impact on HMEC-1 endothelial cells [13,15], we initially sought to determine whether factors present in the CM could modulate Met expression and activation in intestinal tumor cells.

To address this, TCM was generated from HMEC-1 cells previously exposed to PTHrP, as mentioned in Materials and Methods. HCT116 cells were then treated with TCM for different time periods. Met protein expression and activation were subsequently assessed by Western blot analysis using specific antibodies against total Met and activated/phosphorylated Met (Tyr1234/1235). As seen in **Figure 1**, treatment with TCM for 60 minutes increases the protein expression of Met, without substantial modifications in the expression of the phosphorylated protein, compared to the control (CCM). However, p-Met protein levels increased significantly after 5 hours of TCM exposure, compared to the control. We observed that in the condition where the level of p-Met was incremented, the level of total Met decreased (arrows in **Figure 1**), probably because, as happens with other RTKs on the cell surface, upon activation/ phosphorylation Met is internalized for proteasomal degradation [33,34]. In this sense, these findings suggest a modulation of Met expression and activation induced by the action of TCM derived from HMEC-1 endothelial cells exposed to PTHrP.

**Figure 1.**
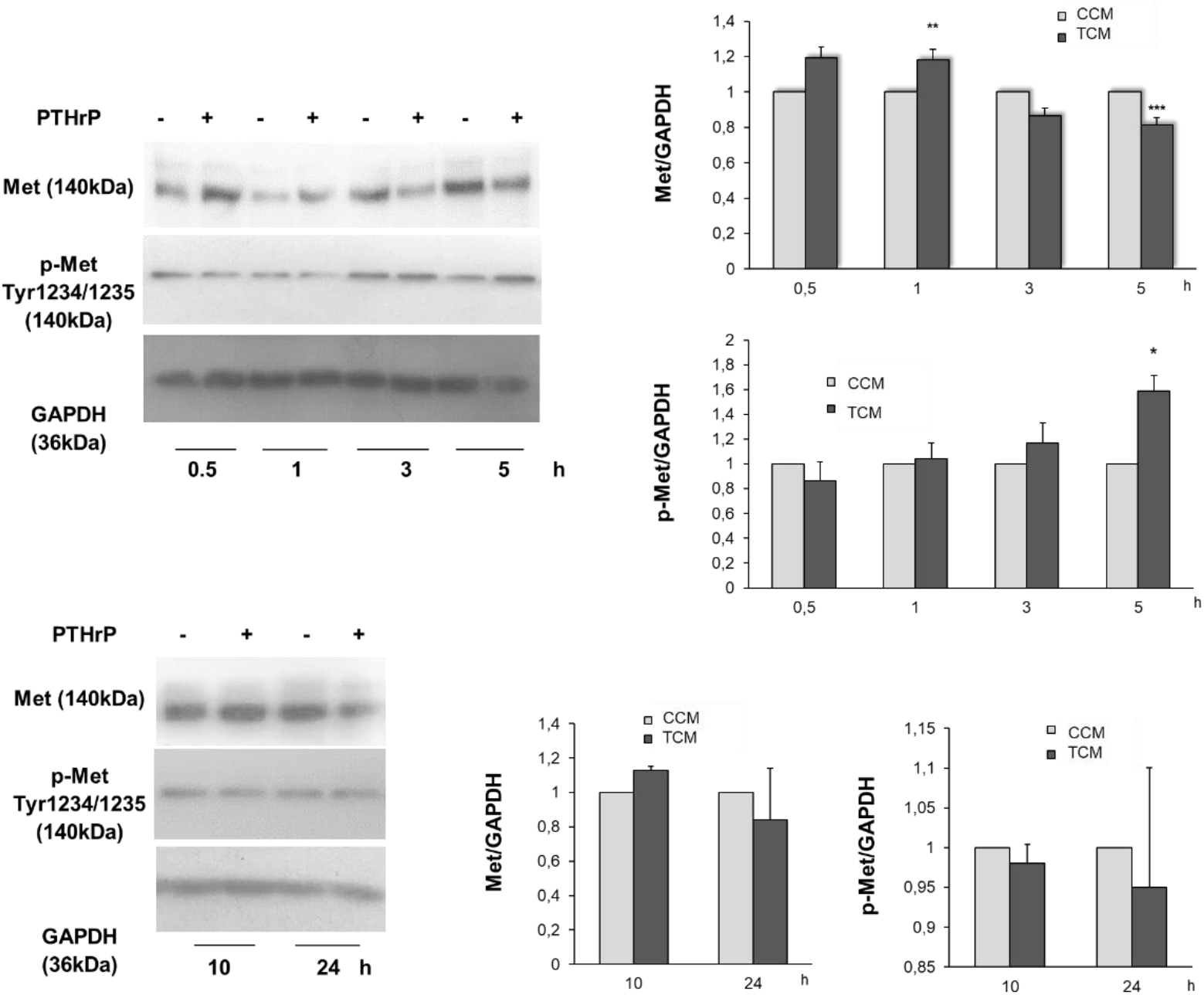
The CM derived from HMEC-1 endothelial cells treated with PTHrP modulates Met protein expression in HCT116 cell line. Cells were treated with CCM or TCM at different times. The protein levels of pro-Met (Met precursor), Met (mature form) and the activated form, p-Met (Tyr1234/1235), were analyzed by western blot to investigate if the CM is capable of modulating the expression and activation of the RTK Met in HCT116 cells. The protein expression levels of GAPDH were determined as a control for the amount of proteins present in the membrane, since this protein is not substantially modified with treatment by the cytokine. Data are presented as mean ± SEM. Statistical analyses were performed on data derived from three independent experiments (n=3). *p<0.05; **p<0.01; *** p<0.001. GAPDH, glyceraldehyde 3-phosphate dehydrogenase; CCM, control conditioned medium; Met, Met receptor tyrosine kinase; p-Met, phospho-Met; PTHrP, parathyroid hormone-related peptide; TCM, treated conditioned medium.

### 2.2. The conditioned media from HMEC-1 cells exposed to PTHrP promotes CSC-associated features in the HCT116 cells

Within the TME, cancer cells exhibit remarkable plasticity, enabling transient and reversible phenotypic transitions that facilitate adaptation to environmental cues. In CRC, disease progression is characterized by key plasticity-related events, including EMT and the acquisition of cancer stem cell CSC traits [35]. In our previous work, we demonstrated that TCM induces phenotypic and molecular changes consistent with EMT in HCT116 cells [13]. Based on these findings, we next investigated whether TCM derived from PTHrP-treated HMEC-1 cells could also promote the acquisition of CSC-associated features. To that end, the protein expression of CD44, a key CSC marker in CRC, was studied. This marker was selected not only by its association with the evolution and poor prognosis of CRC [10,36-38] but, for the close relationship between the expression of Met and CD44, and their functional interaction in this type of tumor [39]. HCT116 cells were exposed to TCM at different times and, by Western blot analysis, a significant increase in CD44 protein expression was observed after 5 hours, compared to CCM (**Figure 2**). Furthermore, the protein levels of this marker were negatively modulated at 24 hours. The observed increase in CD44 protein levels interestingly coincides with Met activation (see black arrows in **Figure 1**).

**Figure 2.**
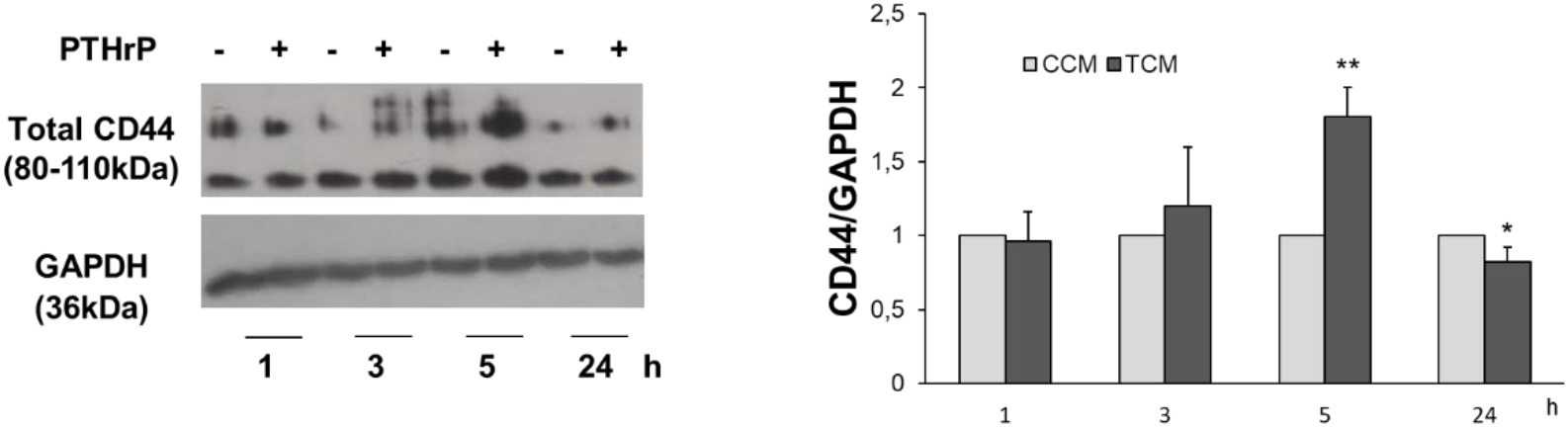
The CM derived from HMEC-1 endothelial cells treated with PTHrP modulates CD44 protein expression in HCT116 cell line. Cells were treated with CCM or TCM at different times. The protein expression of CD44 was analyzed by western blot to investigate if the CM modulates the expression of molecular markers associated with CSC. The protein levels of GAPDH were determined as a control for the amount of proteins present in the membrane since this protein is not substantially modified with the treatment Data are presented as mean ± SEM. Statistical analyses were performed on data derived from three independent experiments (n=3). *p<0.05; **p <0.01. CCM, control conditioned medium; CD44, homing cell adhesion molecule; CSC, cancer stem cells; GAPDH, glyceraldehyde 3-phosphate dehydrogenase; PTHrP, parathyroid hormone-related peptide; SCM, stem cell medium; TCM, treated conditioned medium.

To further evaluate the properties of CRC-derived CSC, their effect on the colonosphere formation capacity was also explored. HCT116 cells were seeded in a monolayer and then exposed to TCM or CCM for 5 hours. This treatment time was selected since a significant increase in CD44 was previously observed at 5 hours of exposure to CM. After this time, the cells were washed, trypsinized and seeded at 10 cells per well in an ultra-low attachment 96-well plate, in SCM. The incubation of the cells was carried out for 12 days and through microscopic observation, the formation of the spheroids was recorded during this period. Representative photographs of each well, taken on day 2, 4, 6, 8, 10 and 12, and the quantification of the results are shown in Figure 3. A significant increase in the area of the colonospheres was detected in cells exposed to TCM compared to area values in control wells (**Figure 3A**). On the other hand, on day 12 of incubation with the TCM, a significant increase was recorded in the number of spheroids formed, concerning the CCM (**Figure 3B**).

**Figure 3.**
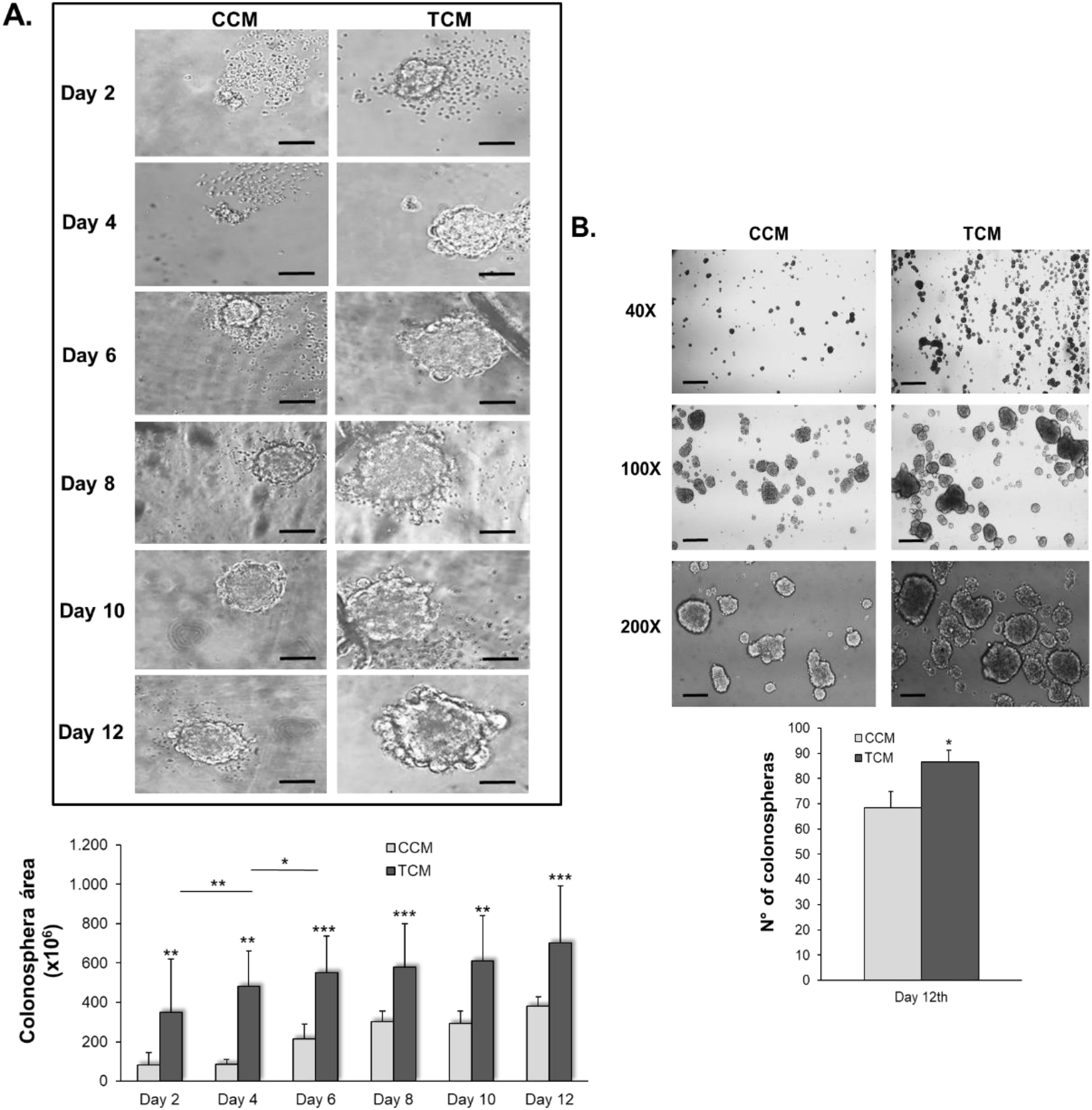
The CM derived from HMEC-1 endothelial cells treated with PTHrP induces an increase in the area and number of colonospheres generated from the HCT116 cell line. HCT116 cells were seeded in a monolayer and treated with CCM or TCM for 5 hours. Cells were subsequently washed, trypsinized and seeded at low density in a non-adherent culture plate in SCM. Representative photographs were taken of each well for 12 days (400x). (A) The analysis of the images with the Image J-NIH program revealed that PTHrP induces an increase in the area of spheroids with respect to controls. (B) At day 12th, PTHrP induces an increase in the number of spheroids with respect to controls. Scale bar, 100um. Data represent the mean ± SEM of two independent experiments (n=2), each performed with four technical replicates per condition. *p<0.05; **p<0.01; ***p<0.001. CCM, control conditioned medium; PTHrP, parathyroid hormone-related peptide; SCM, stem cell medium; TCM, treated conditioned medium.

Collectively, these findings support the hypothesis that soluble mediators present in the TCM may regulate CSC-associated features in HCT116 cells.

### 2.3. The conditioned media from HMEC-1 cells exposed to PTHrP promotes an increase in the number of viable cells and attenuates the effect of CPT-11 on HCT116 cells through the Met pathway

Previously, we demonstrated that TCM induces cell migration and morphological and molecular changes associated with the EMT program in HCT116 cells [13]. Since these events and CSC traits are strictly related to chemoresistance development [40], and considering that in CRC cells, we found that PTHrP induces resistance to CPT-11 through Met signaling [14,32], we proceeded to analyze the potential role of the CM-Met axis in this phenomenon.

Cells were pre-incubated with SU11274, followed by treatment with TCM or CCM. As mentioned in the Materials and Methods section, the dose of SU11274 was chosen according to the literature [41], and the exposure time to TCM was selected according to our previous results [13-15].

Our recent observation revealed by the Trypan blue dye exclusion technique (**Figure 4A**), that the number of viable HCT116 cells increased after 24 hours of exposure to TCM or TCM plus DMSO (vehicle of the inhibitor) compared to their controls. However, this effect was reversed in the presence of Met inhibitor. The same results were observed through the NRU assay (**Figure 4B**).

**Figure 4.**
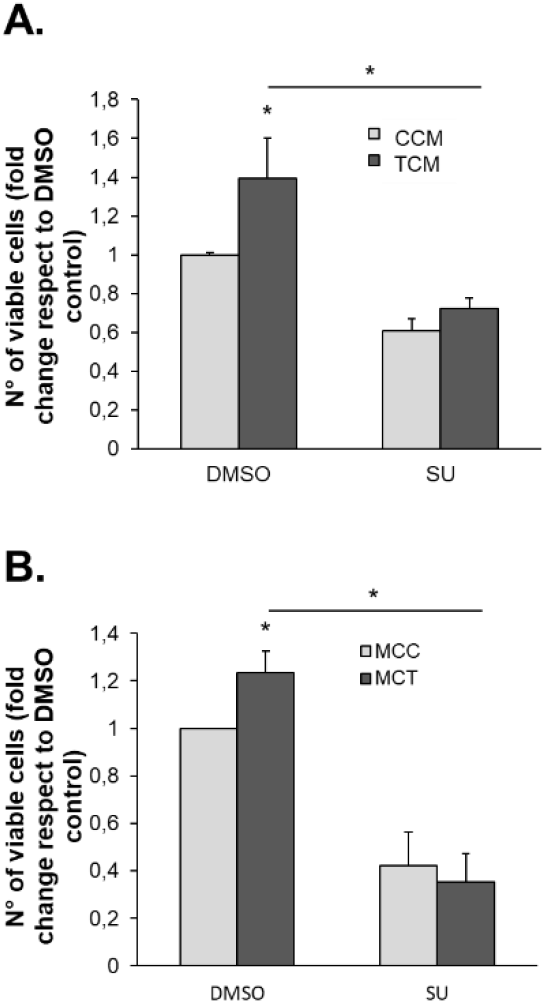
The CM derived from HMEC-1 endothelial cells treated with PTHrP increases the number of viable HCT116 cells through the Met signaling pathway. HCT116 cells were pre-incubated with SU11274 and subsequently treated with CM for 24 hours. The number of viable cells was determined by (A) the Trypan blue test and (B) the NRU assay. In each experiment, a control was performed with DMSO, SU11274 vehicle. Data represent the mean ± SEM of two independent experiments (n=2), each performed with five technical replicates per condition. *p<0.05. CCM, control conditioned medium; DMEM, Dulbecco’s modified Eagle culture medium; NRU, neutral red uptake; PTHrP, parathyroid hormone-related peptide; TCM, treated conditioned medium.

Regarding the evaluation of the effects on the cytotoxicity of the chemotherapeutic drug, the Trypan blue dye exclusion technique (**Figure 5A**) and the NRU assay (**Figure 5B**) revealed that TCM decreases the sensitivity of intestinal tumor cells to CPT-11, through the Met signaling pathway.

**Figure 5.**
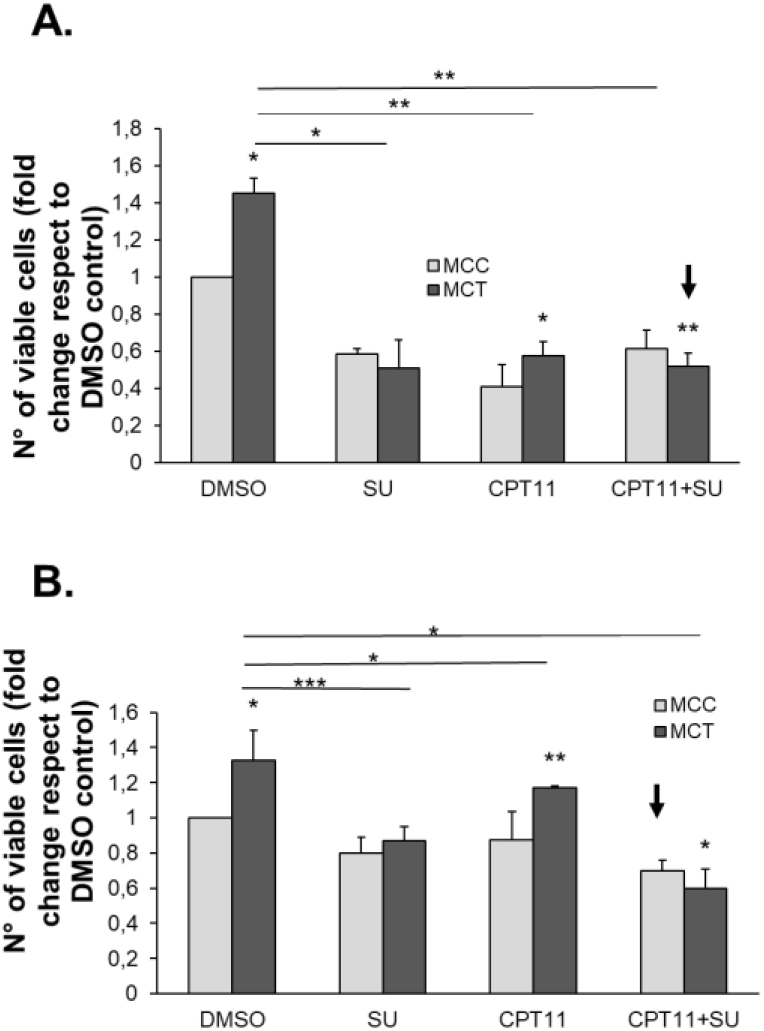
The CM derived from HMEC-1 endothelial cells treated with PTHrP decreases sensitivity to the chemotherapeutic drug CPT-11 in HCT116 cells through the Met signaling pathway. HCT116 cells were pre-incubated with SU11274 and subsequently treated with CM and/or CPT-11 (10 μM) for 24 hours. The number of viable cells was determined by (A) the Trypan blue test and (B) the NRU assay. In each experiment, a control was performed using DMSO as the SU11274 vehicle. Data represent the mean ± SEM of two independent experiments (n=2), each performed with five technical replicates per condition. *p<0.05; **p<0.01; ***p <0.001. CCM, control conditioned medium; CPT-11, irinotecan; DMSO, dimethyl sulfoxide; NRU, neutral red uptake; PTHrP, parathyroid hormone-related peptide; TCM, treated conditioned medium.

These findings suggest that cytokines and/or factors present in the CM derived from HMEC-1 endothelial stromal cells induce an increase in the number of viable cells and attenuate the cytotoxic effect of the drug CPT-11 on HCT116 tumor cells, through the Met signaling.

### 2.4. PTHrP modulates TGF-β1 expression in the HMEC-1 endothelial cell line

As we previously demonstrated, PTHrP increases the expression and release of SPARC, a factor associated with the TME and the aggressive CRC phenotype, in HMEC-1 endothelial cells [13]. Considering the complexity of molecular signals from the microenvironment that favor cancer progression, we next explored whether PTHrP modulates the expression of other factors in endothelial stromal cells that are known to be present in the extracellular environment and participate in this process. The comprehensive molecular characterization performed by Guinney et al. highlighted the critical contribution of TME-related factors to CRC progression, particularly in the CMS4 subtype, which exhibits increased expression of genes associated with stromal activation, including the growth factor TGF-β1 [15]. More recent studies have further demonstrated the overexpression and dysregulation of TGF-β1 signaling across multiple tumor types, including CRC [42,43]. In particular, it has been shown that several TME-derived cell populations, such as cancer-associated fibroblasts (CAF) and endothelial cells, produce and secrete multiple cytokines, among which TGF-β1 plays a central role [4,44,45]. Moreover, CRC-associated endothelial cells display a positive correlation between SPARC and TGF-β1 gene expression compared with endothelial cells derived from healthy intestinal tissue [44].

This evidence motivated us to explore whether PTHrP participates in the modulation of the synthesis and secretion of TGF-β1 in endothelial stromal cells. Based on the previous results, TGF-β1 protein expression was evaluated in the HMEC-1 line. As can be seen in **Figure 6**, through Western blot analysis, it was observed that the TGF-β1 protein levels increased significantly after 16 hours of exposure to PTHrP.

**Figure 6.**
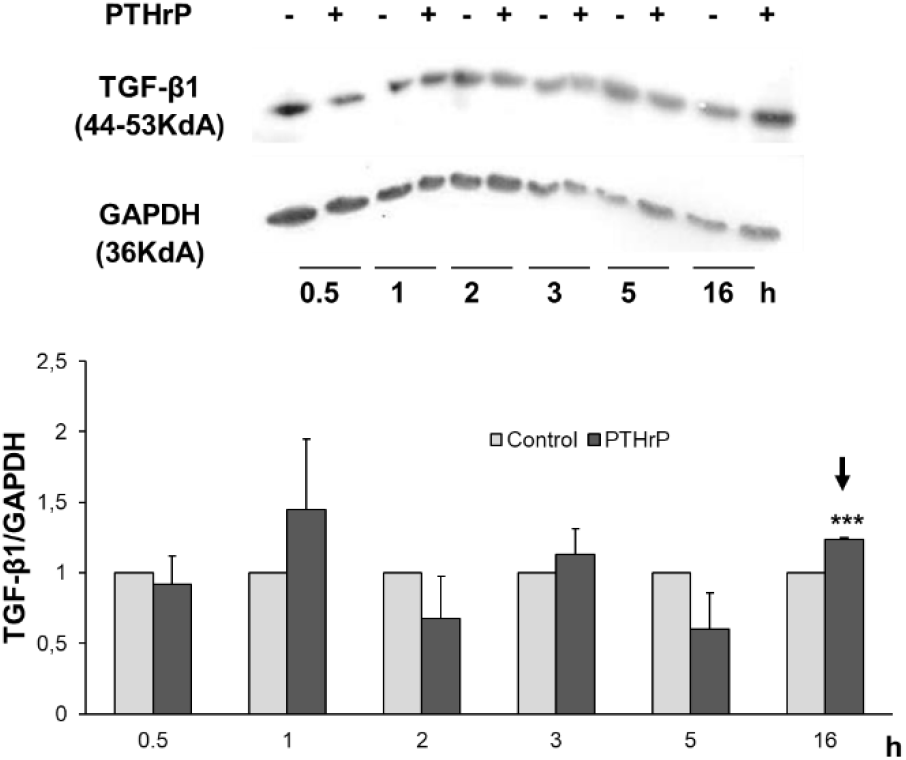
PTHrP modulates TGF-β1 protein expression in HMEC-1 stromal endothelial cells. Cells were treated with or without 10^-8^ mol/L PTHrP at different times. The protein expression of Transforming growth factor beta 1 (TGF-β1) was analyzed by western blot to investigate whether the molecular mechanisms triggered by PTHR1 after PTHrP binding modulate the expression of this factor in HMEC-1 cells. The protein levels of GAPDH were determined as a control for the amount of proteins present in the membrane since this protein is not substantially modified with the treatment. Data are expressed as mean ± SEM. Statistical significance was determined from three independent experiments (n=3).***p<0.001 GAPDH, glyceraldehyde 3-phosphate dehydrogenase; PTHrP, parathyroid hormone-related peptide; TGF-β1, transforming growth factor beta 1.

Although TGF-β intracrine signaling mechanisms have been reported [46], the main mechanism of action of TGF-β1 and other isoforms (TGF-β2 and TGF-β3), is given by its autocrine and paracrine activity [47]. For this reason, after its synthesis and post-translational modification, TGF-β1 is secreted into the extracellular environment [47-49].

Taken together, these data and observations suggest that PTHrP promotes SPARC and TGF-β1 protein expression and secretion in HMEC-1 endothelial stromal cells.

### 2.5. The conditioned media derived from HMEC-1 cells exposed to PTHrP promotes an increase in the number of viable cells and attenuates the cytotoxic effect of the chemotherapeutic drug CPT-11 on the HCT116 cell line, through TGF-β1

TGF-β1 is one of the most relevant growth factors within the TME of CRC [42-45]. As we mentioned above, its physiological activity is subverted during tumor development to promote carcinogenesis, cell proliferation, invasion, migration, a permissive TME with immune evasion, and CRC-associated chemoresistance [4,17,18,50]. Considering the results obtained so far and the role of TGF-β1 in CRC chemoresistance [4,17], we analyzed the effect of this factor, present in the CM, on intestinal tumor cells. For this purpose, we pre-incubated the TCM and CCM with anti-igG or anti-TGF-β1 antibody; then HCT116 cells were exposed to these CM and subsequently treated with the chemotherapeutic drug CPT-11 or its vehicle of dilution. The Trypan blue dye exclusion technique revealed that protein neutralization of TGF-β1 reverses the effect of TCM on the number of viable cells (**Figure 7**, red arrow); however, no statistically significant changes were observed in the resistance to CPT-11, compared to the control (**Figure 7**, green arrow). To corroborate these observations, intestinal tumor cells were directly exposed to TGF-β1 (10ng/ml). This concentration was selected according to previous literature [51,52]. The Trypan blue dye exclusion assay shows that this growth factor promotes an increase in the number of viable cells, without inducing statistically significant changes in their sensitivity to CPT-11 **(Figure 8**, red arrows). In parallel, it was evaluated whether the combined treatment of HCT116 cells with TGF-β1 and PTHrP promoted a synergistic or antagonistic effect on the action of these molecules. As seen in **Figure 8 (**green arrows), the activity of TGF-β1 does not antagonize or synergizes the effect of PTHrP on the number of viable cells or on the sensitivity to the chemotherapeutic drug studied. These findings suggest that TGF-β1 produced by endothelial stromal cells HMEC-1 in response to PTHrP, and present in the CM, participates in the promotion of events associated with CRC progression. Furthermore, the activity of this factor seems not to interfere with the effect of PTHrP on intestinal tumor cells.

**Figure 7.**
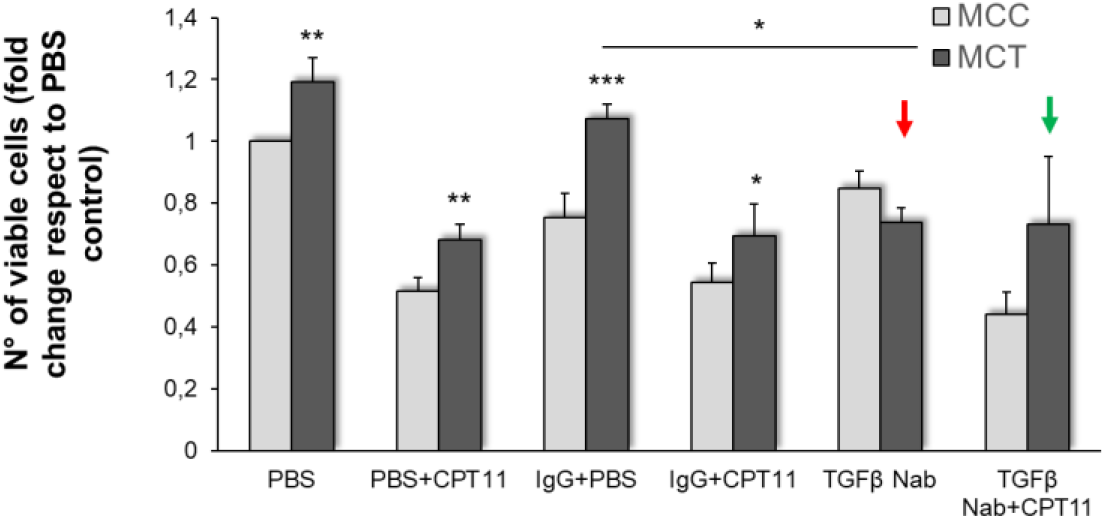
The CM derived from PTHrP-treated HMEC-1 stromal endothelial cells increases the number of viable HCT116 cells through the TGF-β1 signaling pathway. Cells were exposed to CM preincubated with PBS or anti-IgG or anti-TGFβ1 and then treated with CPT-11 or its vehicle (DMSO). The number of viable cells was quantified using the trypan blue technique. Data are presented as mean ± SEM. Statistical analyses were performed on data derived from three independent experiments (n=3), three technical replicates each. *<0.05; **p <0.01; ***p<0.001. CCM, control conditioned medium; CPT-11, irinotecan; DMSO, dimethyl sulfoxide; PTHrP, parathyroid hormone-related peptide; TCM, treated conditioned medium; TGF-β1, transforming growth factor beta 1.

**Figure 8.**
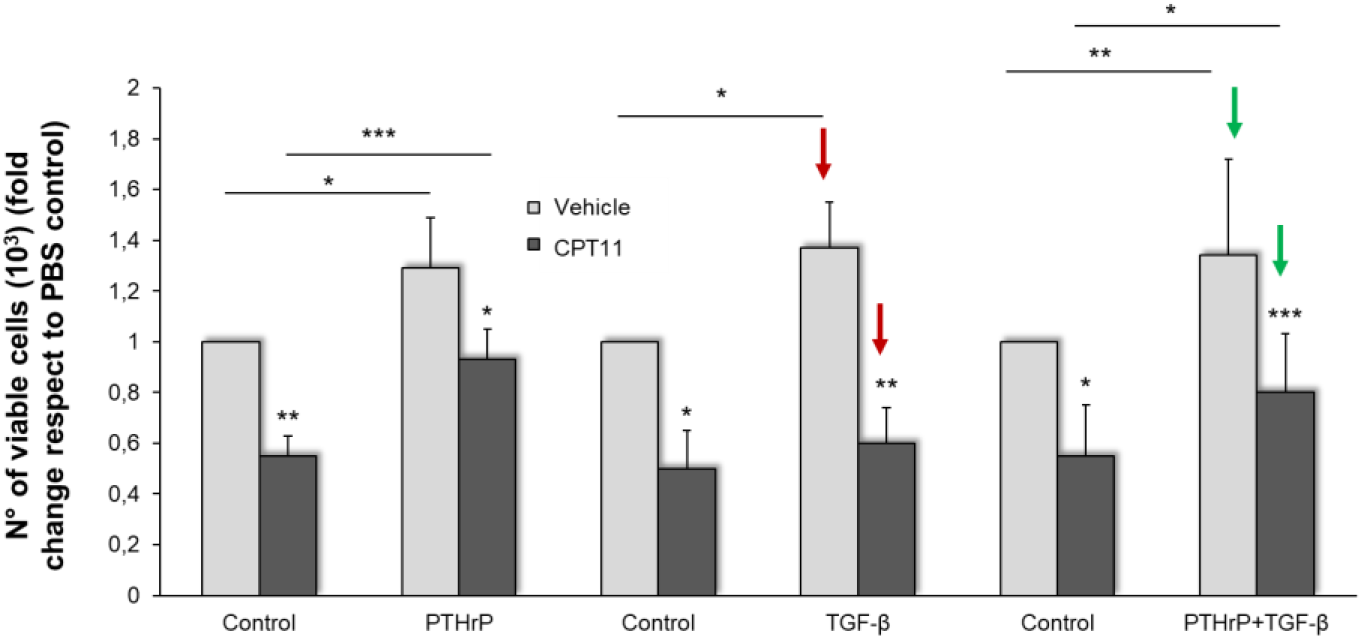
Treatment with PTHrP and/or TGF-β1 promotes an increase in the number of viable cells in HCT116 cells. Cells were treated with PTHrP, TGF-β1, or both cytokines together, and then exposed to CPT-11 or its vehicle. The number of viable cells was quantified using the Trypan blue technique. Data are presented as mean ± SEM. Statistical analyses were performed on data derived from two independent experiments (n=2), three technical replicates each. *p<0.05; **p <0.01; ***p <0.001. CPT-11, irinotecan; PTHrP, parathyroid hormone-related peptide; TGF-β1, transforming growth factor beta 1.

### 2.6. Analysis by exploring databases of genes expressed in the tumor microenvironment of patients with CRC

As seen in **Figure 9**, a GEO2R analysis was performed on samples of CRC stroma compared to healthy patients’ intestinal stromal tissue (Accession number: GSE31279). The list of overexpressed and downregulated DEGs in the data set (Log_2_FC>1 or Log_2_FC<-1 and Benjamini and Hochberg adjusted p < 0.05) are shown in **Supplementary material, Table 1**. The DEGs resulting from this analysis were incorporated into the Cytoscape 3.10.0 software to create an interaction network and find the most significant functional interaction module according to the selected criteria (bibliographic data and experimental results). We observed that the protein induced by transforming growth factor beta (TGFβI), which is regulated by TGF-β signaling and whose increased expression is related to a poor prognosis in CRC [53], is overexpressed in this module (black arrow in **Figure 10**).

**Figure 9.**
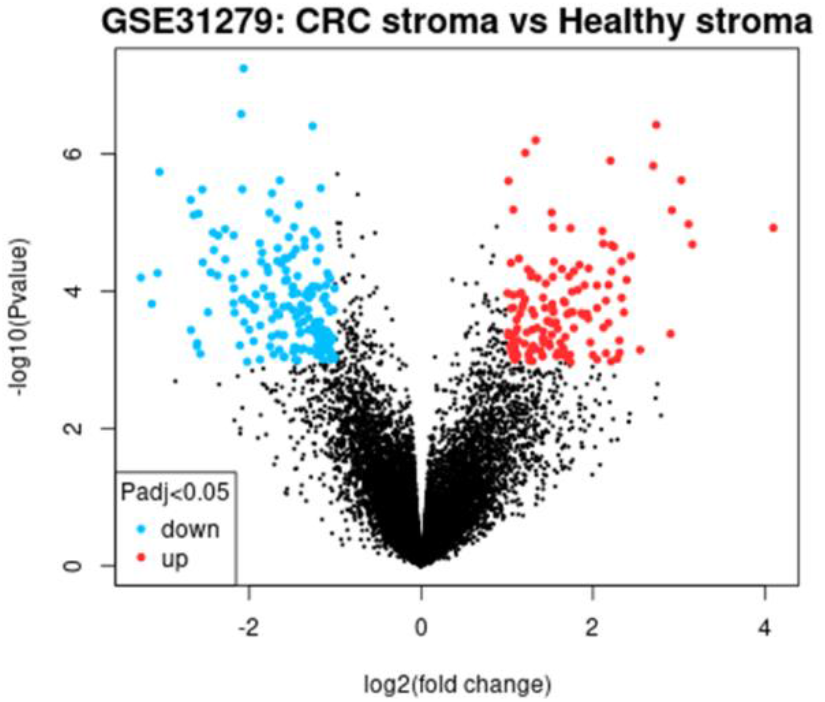
Differentially expressed genes in intestinal stromal tissue of patients with CRC with respect to healthy stromal tissue. Volcano diagram. Differentially expressed genes in CRC stromal tissue were obtained by GEO2R (red: over-expressed; light blue: under-expressed). P adjusted by Benjamini and Hoechberg <0,05; Fold change 1 and -1.

**Table 1.** Clinicopathological characteristics of the patients.

| Variable | Value |
| --- | --- |
| Sex *n* (%) |  |
| Female | 6 (55%) |
| Male | 5 (45%) |
| Age median (range) | 61 |
| Primary tumor site *n* (%) |  |
| Right-side | 4 (36%) |
| Left-side | 3 (28%) |
| Rectum | 4 (36%) |
| Tumor differentiation *n* (%) |  |
| Well differentiated | 6 (55%) |
| Moderately or poorly differentiated | 5 (45%) |
| Initial stage *n* (%) |  |
| I/II | 8 (73%) |
| III | 3 (27%) |

**Figure 10.**
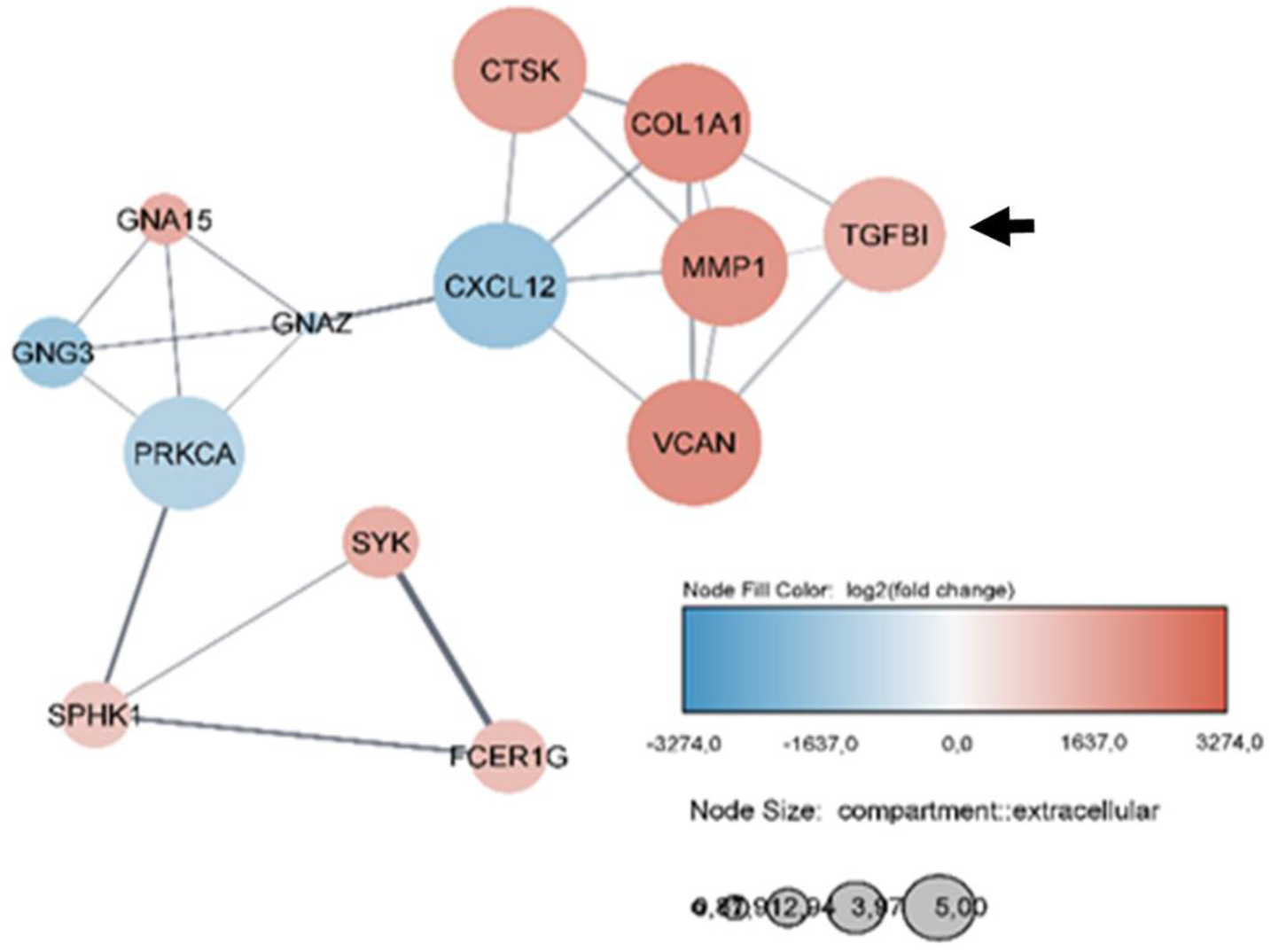
Analysis of differentially expressed genes in intestinal stromal tissue of patients with CRC concerning healthy stromal tissue. Differentially expressed genes obtained by GEO2R were visualized with Cytoscape 3.10.0 software to create an interaction network. The most relevant functional interaction module (based on bibliographic references and experimental data) is constituted by 13 of the 319 differentially expressed genes. The color of the nodes indicates if they are over-expressed (red) or under-expressed (light blue) and their size, the degree of their expression in the extracellular compartment. The intensity and thickness of the axes indicate the strength of the interrelationship between genes.

To delve deeper into the function of the DEGs, their participation in biological processes was evaluated, as well as the signaling pathways in which they are involved. For this, the DEGs were incorporated into the EnrichR-Appyter platform [54]. As seen in **Figure 11**, using the public databases Reactome 2022 and BioPlanet 2019, we observed that the genes overexpressed in CRC are significantly associated with extracellular matrix organization processes (p< 2.87×10^-5^), pathways of signal transduction (p<2.9×10^-3^), Met activation (p< 4.27×10^-4^), and effects triggered by TGF-β (p< 2.14×10^-6^), among others. On the other hand, genes downregulated in samples derived from CRC stroma showed an association with events related to TGF-β (p<1.59×10^-5^) and the modulation of signaling pathways previously studied by us [19].

**Figure 11.**
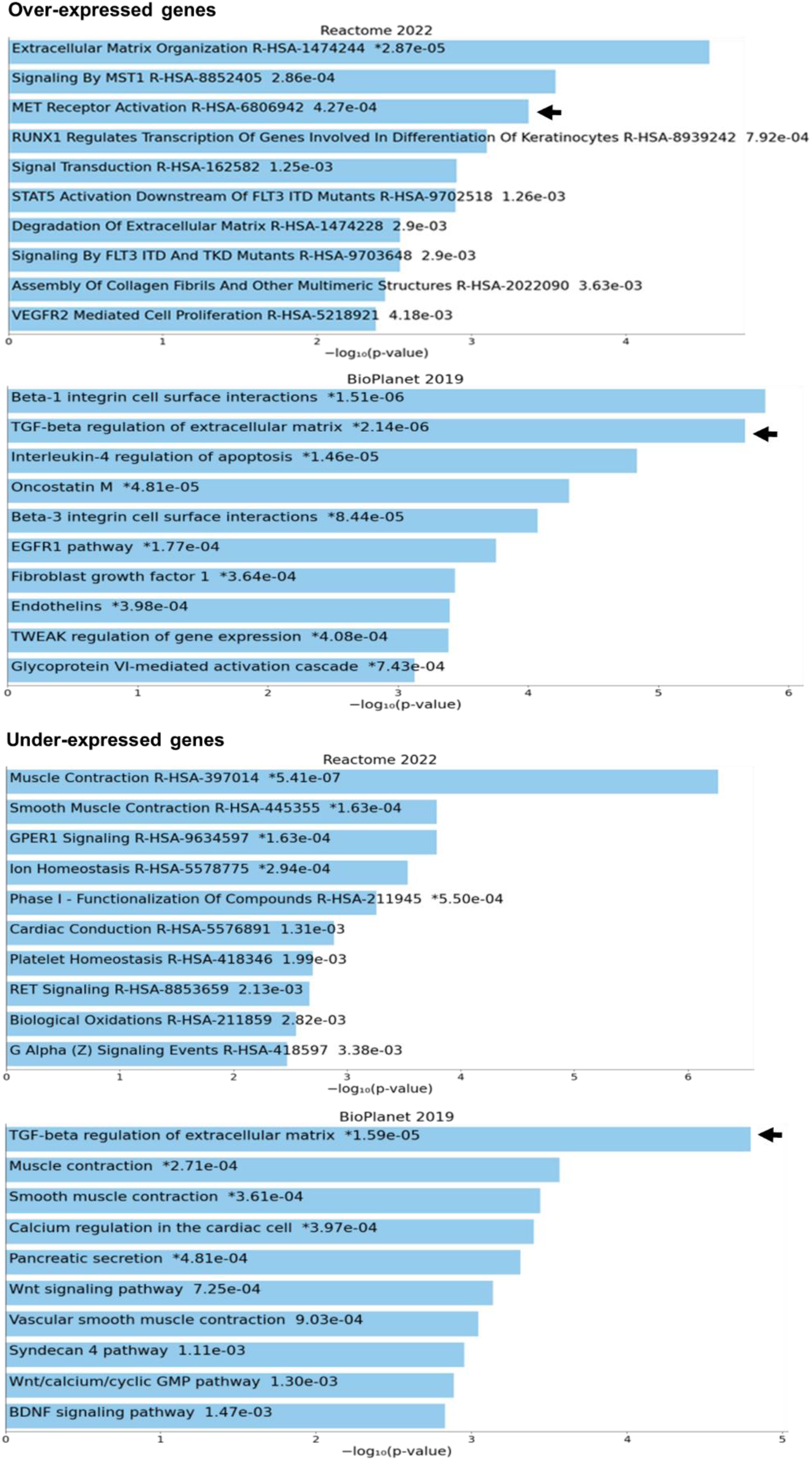
Functional analyses of differentially expressed genes in intestinal stromal tissue of patients with CRC concerning healthy stromal tissue. Using the EnrichR-Appyter platform, the functional enrichment query was carried out by biological processes and signaling pathways, for the over-expressed genes (upper panel) and the under-expressed genes (lower panel). The graph bars show the associated probability to each function expressed as – log10 (p-value <0.05).

### 2.7. Database analysis of the clinical implications of TME markers modulated by PTHrP in CRC

To determine in clinical samples if there is an association between PTHrP and the markers previously evaluated, an analysis was carried out using the Pearson correlation test (R) in paired samples of colon cancer (COAD) and rectal cancer (READ), using the online database GEPIA2. As can be seen in **Figure 12A**, the expression of the PTHrP gene (PTHLH) positively correlated with the gene expression of its receptor PTHR1, TGFBR1 and TGF-β1, although the strength of the association is modest (R=0.22, R=0.39 and R=0.45, respectively) reaches high statistical significance (p=2.4×10^-5^, p=9.8×10^-15^, and p=0, respectively); in line with these observations the same results were seen between the genes encoding PTHR1 and TGF-β1 (R=0.35; p=2.4×10^-12^) or PTHR1 and TGFBR1 (R=0.28; p=3.7×10^-8^). Furthermore, a positive correlation was also observed between TGF-β1 and TGFBR1 genes (R=0.57; p=0). On the other hand, in **Figure 12B**, it is possible to visualize the link between factors that are known to be present in the TME of CRC. A positive and significant correlation was observed between the genes encoding HGF and TGF-β1 (R=0.48; p=0), HGF and SPARC (R=0.54; p=0), or SPARC and TGF-β1 (R= 0.71; p=0).

**Figure 12.**
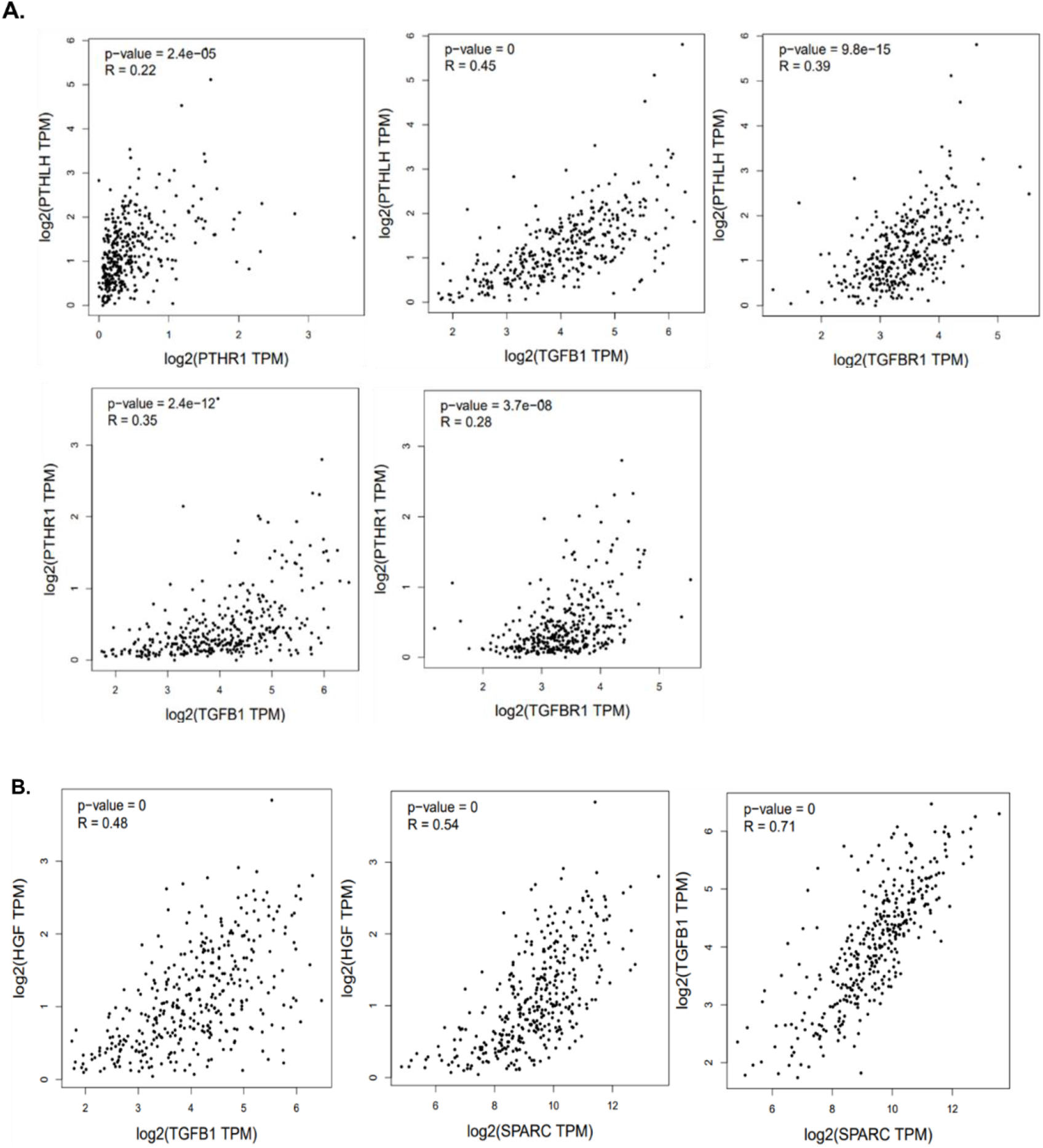
Correlation between the expression of markers associated with CRC. The graphs show, **(A)** the correlation of gene expression between PTHrP (PTHLH) with its receptor (PTHR1), with TGF-β1 (TGFB1) and its receptor (TGFBR1). Additionally, the correlation between both receptors PTHR1 and TGFB1 or TGFBR1; finally, the correlation among TGFB1 and TGFBR1. **(B)** the correlation of HGF gene expression with TGF-β1 and with SPARC, and of SPARC with TGF-β1, using the GEPIA2 platform that analyzes data from CRC tumor samples from the TCGA repository. CRC, colorectal cancer; PTHrP, parathyroid hormone-related peptide; HGF, hepatocyte growth factor; SPARC, secreted protein acidic and cysteine-rich; TGF-β1, transforming growth factor beta 1.

All these findings reinforce the hypothesis of the participation at least of PTHrP, TGF-β and Met signaling pathways in the tumor bulk of CRC patients.

### 2.8 Evaluation of TGFBR1 and PTHR1 expressions in CRC human samples

The findings presented in previous sections, derived from *in vivo* assays, indicate that the activation of PTHR1 following its interaction with PTHrP can induce the expression and secretion of TGF-β1 by stromal endothelial cells. Our observations suggest that this factor mediates effects in CRC-derived cells that are promoted by TCM. Additionally, our *in silico* analyses we analysed the correlation among gene expression of PTHrP, TGF-β1, their receptors and associated factors within human CRC samples. Given the statistical significance of the data and the notable trend towards a positive correlation among these molecules, we aimed to further clarify these associations. Since in our previous work we found that PTHR1 expression was statistically lower in tumors with moderately or poor histological differentiation [4], we decided to confirm the behavior of PTHR1 expression and complemented this information with TGFβR1 expression assessment in human CRC samples to ascertain potential associations with tumor characteristics. A total of 11 samples from patients diagnosed with colorectal adenocarcinoma, along with 2 samples of normal colonic mucosa, were analyzed through immunohistochemistry. The characteristics of the patient cohort are detailed in **Table 2**. The average age was 61 years; 45% (n = 5) were male, and 55% (n = 6) were female. Concerning primary tumor location, 36% (n = 4) presented right-colon cancer, 28% (n = 3) left-colon cancer, and 36% (n = 4) in the rectum; 55% (n = 6) presented histological well differentiated neoplasia, while 45% (n = 5) moderately or poorly differentiated; 73% (n = 8) of the CRC patients presented stage I or II, and 27% (n = 3) stage III. **Figure 13** shows a comparison of immunostaining for both receptors in healthy tissue and tumors across different histological grades. Both receptors were detected in normal colonic mucosa, consistent with their established roles in intestinal physiological processes. In CRC samples, qualitative and quantitative examination suggested differences in staining intensity among normal tissue and tumor, and also between histological grades. The results suggest that both PTHR1 and TGFβR1 increase their expression in GH1-positive tumors compared to normal tissue. However, in neoplastic tissues, both markers decrease with histological differentiation. Although these observations are descriptive in nature due to the limited number of samples analyzed, they are consistent with our *in vitro* and *in silico* findings and provide additional morphological support for the proposed biological association.

**Figure 13.**
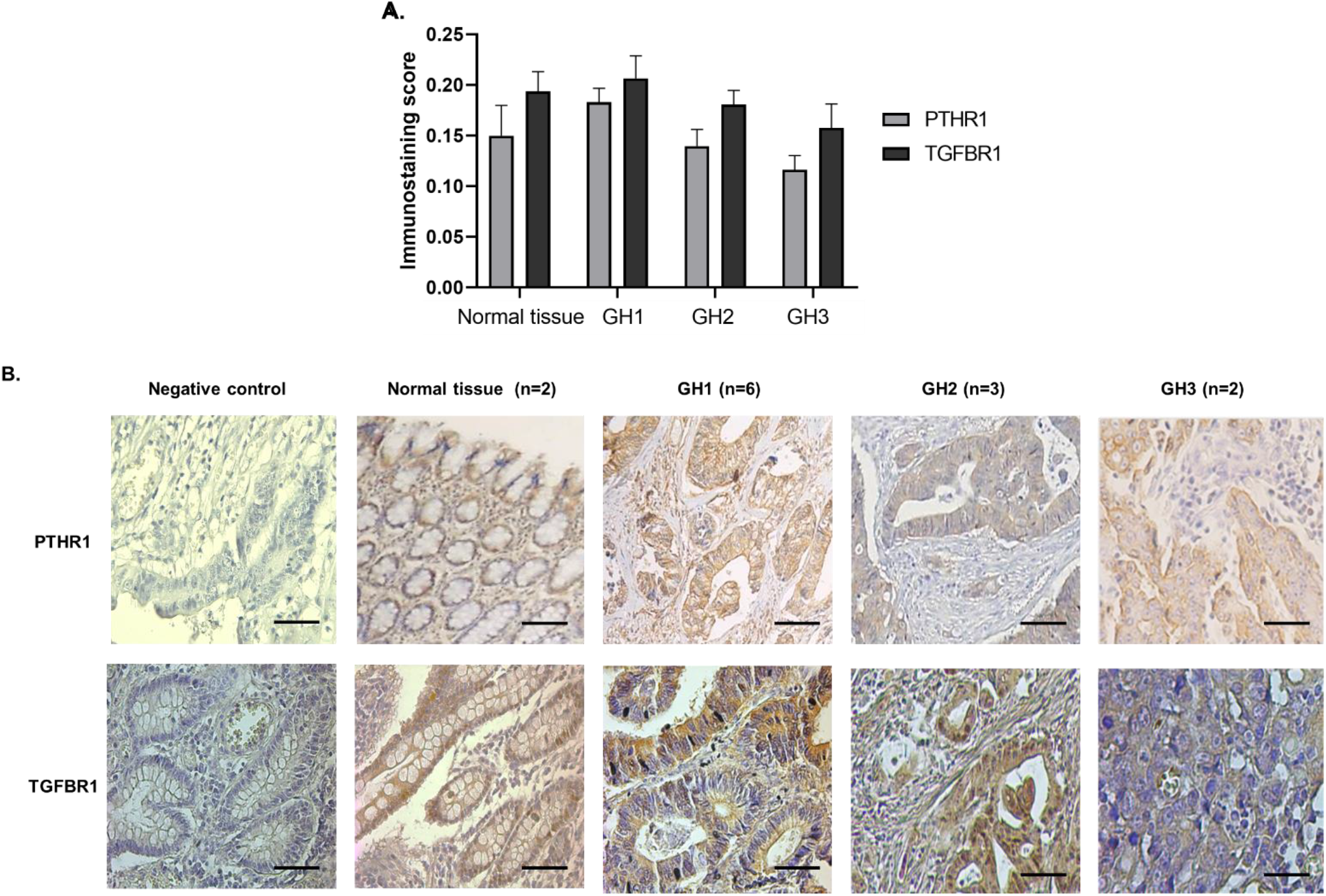
Expression of PTHR1 and TGFBR1 in human samples. Representative images obtained using the IHC technique in normal colon mucosa and tumor tissue with histological grade GH1, GH2 and GH3. Scale bar, 50um.

In addition, using GEPIA2 we analyzed Kaplan-Meier curves of patients with primary CRC tumors and signatures of overexpressed genes. As indicated by the Logrank test [55], and the Cox regression test with a hazard ratio (HR) value [56], a downward trend was observed in the overall survival between those patients with or without overexpression of the genetic signature PTHR1/TGFBR1, HR=1.5 Logrank p=0.065 (**Figure 14A**). However, patients with overexpression of PTHR1, HR=1.2 Logrank p=0.34 (**Figure 14B**) or TGFBR1, HR=1.1 Logrank p=0.61 (**Figure 14C**) alone do not show a decreased overall survival compared to those who do not overexpress these genes. Furthermore, the analysis of the disease free survival curves indicates with high probability that those patients with this overexpressed signature have almost double risk of a recurrence compared to patients who have a low expression of this set of genes, HR=1.6; p=0.032.

**Figure 14.**
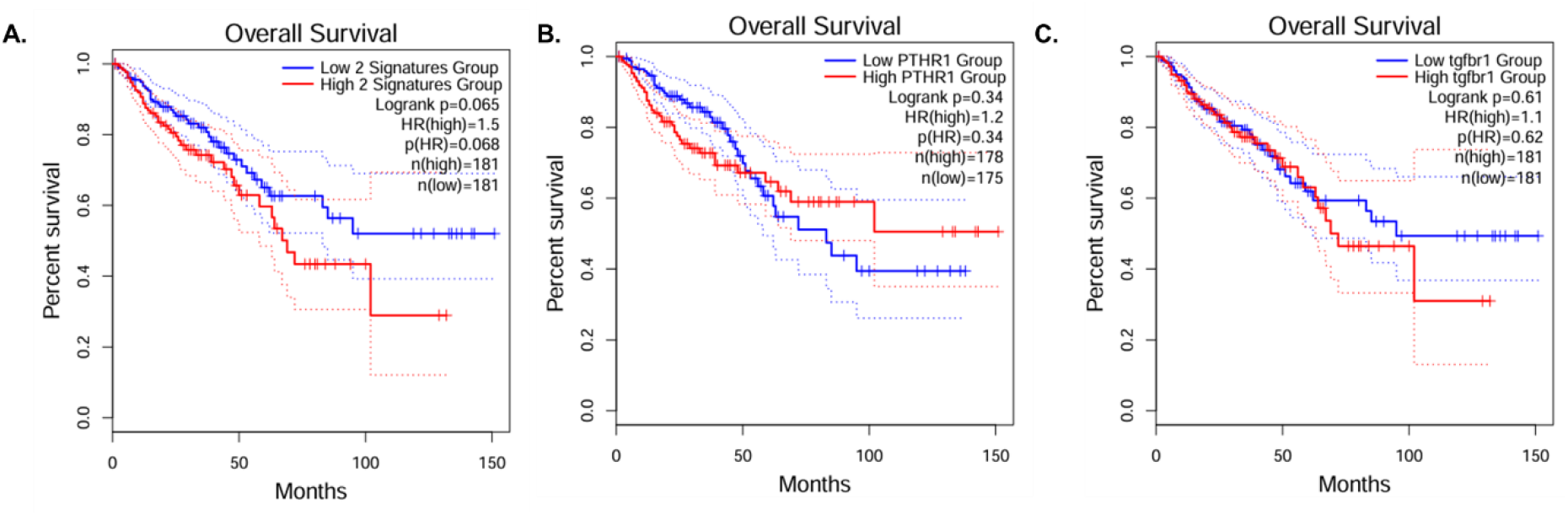
Analysis of the expression of PTHR1 and TGFBR1 in the evolution of patients with colorectal cancer. (A) Curves for overall survival regarding the genetic expression levels of (A) PTHR1/TGFBR1, (B) PTHR1 and C) TGFBR1. Information is presented as Kaplan-Meier plots with data from the Cancer Genome Atlas (TCGA). With a solid red curve, cases with high levels of standardized expression of both markers are shown in each graph, while those with low levels are shown in blue. The curves corresponding to the 95% confidence interval are plotted as dotted lines. The curves are compared using the Logrank (Mantel-Cox) test and its probability value as a potential biomarker in associated cancer (p). The Hazard Ratio (HR) and its respective p are detailed, comparing the high and low expression of the marker groups.

The prognostic impact appears to depend on the co-activation of the pathway, rather than on the overexpression of each gene individually. The combined activation of these receptors could be associated with tumor progression or relapse, even if the impact on overall survival is more moderate.

## 3. Discussion

Recent research indicates that tumors develop and evolve through complex interactions with the surrounding stroma, in addition to simple isolated genetic and epigenetic alterations [57,58]. The TME and the signaling pathways associated, play a crucial role in tumor proliferation, invasion, metastasis, and chemoresistance [4,59,60]. Statistics indicate that around 20% of advanced CRC patients exhibit resistance to commonly used treatments, presenting relapses and poor overall survival [4,61]. In this context, it became necessary to expand knowledge about the mechanisms involved in CRC treatment failure. Regarding this fact, the TME became the focus of these investigations and an unquestionable target for the development of novel therapies [4,57]. Thus, based on these bibliographic data and our previous results, we began to investigate the role of the TME in our experimental model.

Given that in previous works it had been observed that the CM derived from HMEC-1 cells treated with PTHrP promote phenotypic and molecular changes associated with TME and invasion in CRC cells; and that this phenomenon induced by the direct action of the cytokine was mediated by the RTK Met, we began studying whether TCM modulates the expression and activation of this receptor on intestinal tumor cells. As expected, TCM modulates Met protein expression and activation. As it was explained before, the decrease in Met protein expression, simultaneously with an increase in its active/phosphorylated form (Tyr1234/1235) could be due to a degradation of the RTK after its activation [33,34]. This result suggests that the HGF could be present in the TCM, activating the RTK directly, as it is the only known ligand of this receptor. Although there is currently no scientific evidence of the HGF synthesis and release by tumor stromal endothelial cells, it is known that these cells are one of the primary sources of CAF [62], and it is well documented that CAF are important HGF producers in the TME [4,63,64]. In line with these data, it is possible to hypothesize that PTHrP could induce characteristics in endothelial cells that favor the synthesis and secretion of HGF. On the other hand, it is necessary to consider that PTHrP could promote and modulate its own expression in HMEC-1, and in this case, this cytokine present in the TCM would be responsible for the synthesis and activation of Met, as we found in previous works [32]. HMEC-1 expresses PTHR1 [15], although the intrinsic production of PTHrP in this cell line has not yet been elucidated, its synthesis and release have been reported in human endothelial cells [65]. Regarding this, more studies are necessary to identify the presence of these factors in the TCM.

Initially, the expression of the molecular marker associated with CSC, CD44, was analyzed in intestinal tumor cells in response to exposure to CM derived from HMEC-1. Our findings suggest that TCM acts on CRC-derived cells by modulating the expression of CSC markers such as CD44 through Met signaling. However, more studies are necessary to confirm this mechanism and whether TCM is capable of also modulating the expression of other surface markers associated with this cell population.

As previously mentioned, the selection of CD44, CSC marker, is based on its association with the evolution and poor prognosis of CRC [10,36-38]. Additionally, there exists a close relationship between the expression of Met and CD44, which functionally interact with each other in this type of tumor [39]. In fact, the increase in the protein expression of CD44 in HCT116 observed after 5 hours of exposure to TCM, concur with the activation of Met in our experimental system. Similar results have been obtained by other research groups. Wielenga and colleagues reported that CD44 molecules associated with heparan sulfate chains (CD44-HS) bind to HGF on the cell surface and strongly promote Met signaling [66]. Elliott and collaborators found in an *in vivo* model that the activation of Met receptor in intestinal tumor cells is associated with the overexpression of CD44 [67]. Joosten showed in his recent publication that Met activity in intestinal tumorigenesis is promoted by different CD44 isoforms expressed by stem cells. Furthermore, it explains that Met has a similar role to the endothelial growth factor receptor (EGFR), a usual target of immunotherapies in intestinal tumors, and that CRC cells can co-opt the HGF/Met/CD44 pathway to escape therapies directed at EGFR [39]. Given that the TCM and its factors induce Met activation and increase CD44 expression, these events could be associated. Further studies are needed to elucidate the underlying mechanisms. Conversely, a decrease in CD44 expression was observed in intestinal tumor cells after 24 hours of treatment with the TCM, as confirmed by Western blot. This could be linked to the phenomenon of cellular plasticity. Our previous studies have shown that EMT-like phenotypic changes occur in HCT116 cells after 24 hours of exposure to TCM [13]. Therefore, the decrease in CD44 protein expression may be associated with a switch to a mesenchymal phenotype in the cells.

Regarding CSC, we also evaluated the ability of TCM to form colonospheres under conditions that favors the reprogramming/dedifferentiation of epithelial cells into stem cells. As mentioned in Materials and Methods section, the use of ultra-low adhesion culture dishes and a specific serum-free medium is a widely recognized technique to promote the enrichment of the culture in cells with CSC characteristics [68-71]. In line with the results presented before, the colonosphere assay allowed us to observe that TCM induces an increase in the size and number of the spheroids, with respect to the control. Together these observations suggest that TCM fosters traits associated with this cellular subpopulation. This hypothesis is supported by previous research conducted by Wei and collaborators, who in the last decade demonstrated that endothelial cells and/or their precursors can increase the tumorigenic capacity, a characteristic associated with CSC, in CRC derived cells [71]. There is plenty of information on the role of CSC in angiogenesis and neovascularization [72,73], however there is limited knowledge to date on how the collaboration of endothelial cells within the tumor niche can contribute to the induction and sustenance of CSC. It is clear that in the TME, cells have diverse functions. Endothelial cells play a role in the formation and maintenance of vasculature, but also influence the phenotype of CSC by secreting soluble factors. Dr. Ellis’ research group discovered that vascular endothelial cells release Jagged-1 factor activating signaling pathways in colorectal CSC and contributing to their maintenance and survival [74,75]. These data support our results and reinforce the intention to deepen the study of the factors present in TCM.

Given that the biomolecules present in TCM evidently promote characteristics associated with cellular plasticity, dedifferentiation, and resistance to treatment, the effect of TCM on the cell viability and chemoresistance to CPT-11 was investigated. We observed that Met activation mediates an increase in the number of viable cells in HCT116 cells induced by TCM. Furthermore, the exposure of CRC-derived cells to this medium decreases their sensitivity to the chemotherapeutic drug CPT-11, through the Met signaling pathway. These observations were in accordance with other investigations that highlight the relevance of the HGF/Met axis in the TME, as a mediator of the proliferation of intestinal tumor cells [76,77]. However, more studies are necessary to confirm whether the increase in the number of viable cells is due to an increase in survival or an induction of cell proliferation in our model.

Regarding chemotherapeutic resistance, endothelial cells are linked through three interconnected pathways: the first is angiogenesis and vascularization, which allows tumor perfusion and regulates the arrival of chemotherapeutic drugs and the effectiveness of radiotherapy [78]; the second pathway is associated with factors released by both endothelial cells and tumor cells in response to hypoxia, caused by the formation of tortuous and disorganized tubes that do not allow adequate irrigation [15,79,80]; finally, the interaction of endothelial cells with other cell types in the TME [81]. Therefore, the exploration of each of these aspects in our system deserves greater attention.

TGF-β is one of the most studied factors in the tumor stroma to date, and is synthesized and secreted predominantly by endothelial cells and fibroblasts [4,18,72]. It has been established that this factor is associated with several events in the development and progression of CRC. These events include proliferation [82], chemoresistance [83], as well as with the EMT, invasion, and migration, which are events involved in the metastatic process [84,85]. Considering the effect of CM on CRC-derived cells and the confirmation of the expression of TGF-β1 receptor in HCT116 cells by Moez and his research team [86], the increase in the protein expression of this factor in HMEC-1 cells due to PTHrP treatment has strengthened the need to investigate its presence in the TCM. Furthermore, the interrelationship between PTHrP and TGF-β1 in the TME had already been reported in several neoplastic models [16,17,20,21]. One of the most studied events in this regard is the positive feedback mechanism that exists between these two molecules in osteolytic metastases of breast cancer [16,87,88].

To evaluate the presence of TGF-β1 in the TCM, a specific antibody was used to block the cytokine’s binding capacity to its receptor, creating steric hindrance. Then, to investigate whether this factor mediates the observed effects, it was studied its effect on the number of viable cells and the phenomenon of chemoresistance which links all the others. No statistically significant changes were detected in terms of chemoresistance to CPT-11, probably this TCM-induced phenomenon is mediated by factors independent of the TGF-β pathway in our system. Also we observed that the number of CRC viable cells increases with TCM treatment, and that this effect is reversed by blocking TGF-β1. This confirms the presence of the growth factor in the TCM. Additionally, we treated the intestinal tumor cells with the growth factor and found that it participates in the modulation of the number of viable cells. Despite literature reporting an intracrine action of TGF-β1, in most cases, the protein expression is directly associated with its secretion and autocrine and paracrine activity [47]. TGF-β1 is produced and released into the extracellular environment by a great diversity of cells in the tumor microenvironment [4]. Since this cytokine is broadly pleiotropic, its synthesis and activation are strictly controlled. This factor is initially translated as inactive proTGF-β1, forms homodimers, and is cleaved to generate the propeptide and the mature chain. In most cells, TGFβ-1 is secreted and stored in the extracellular matrix-bound to latent transforming growth factor β binding proteins [47-49]. An acidic microenvironment, the action of tensile forces within the extracellular matrix and the interaction of this latent complex with fibrins, integrins, and other proteins constitute the major local regulators of TGF-β1 activation in tissues [47]. However, Fridich and collaborators detected that there are also soluble factors present in the TME that allow the activation of TGF-β1 and its binding to the TGFBR2 receptor promoting its dimerization with TGFBR1 [47]. Furthermore, compounds known to be secreted by endothelial cells, such as nitric oxide (NO), also promote the activation of TGF-β1 [89]. On the other hand, exposure of CRC-derived cells together to PTHrP and TGF-β1 did not induce significant changes with respect to the treatment of each of the cytokines separately. These results suggest that the factors studied do not present an antagonistic or synergistic effect in our *in vitro* model, at least in the events evaluated. However, more studies are necessary in order to deepen the functional link between these molecules in CRC.

The results presented in this work raise the question of the interrelation of the Met and TGF-β1 signaling pathways in our system. Breuning and collaborators demonstrated with *in vitro* and *in silico* models of breast cancer that TGF-β1 modulates the expression and activity of Met through two different mechanisms [90]. The first mechanism describes TGF-β1 as an inducer of the expression of the transcription factor C-ets-1, which mediates the overexpression of Met [90-92]. Furthermore, overexpression of C-ets-1 has been shown to strongly promote the EMT program in CRC cells [93]. The second mechanism of Met regulation by TGFβ-1 postulated by Breuning’s group is related to the activity of the microRNA, miR-128-3p, that is negatively modulated by TGFβ-1 and has Met as a direct target. Furthermore, they observed that the decrease in the expression and activity of Met, in this context, directly impairs cell migration induced by the Met signaling pathway [90]. These mechanisms could explain what would be happening in our system. It is possible that PTHrP released by cancer or stromal cells acts through autocrine/paracrine signaling, promotes the synthesis and secretion of TGF-β1 and other factors that favor an aggressive behavior in tumor cells mediated by Met signaling. Given the relevance of both markers in the development and progression of CRC, as well as in the prognosis of these patients, the interrelationship of Met, TGF-β1 and other factors associated with the TME, such as PTHrP, deserves further exploration.

The results indicate the involvement of both, Met and TGF-β1 in the increase in cell viability induced by the TCM. However, neutralization of TGF-β1 present in this medium does not significantly increase sensitivity to the chemotherapeutic drug CPT-11, unlike what was observed after Met inhibition. Considering that the TCM likely contains multiple soluble factors, both Met and TGF-β could converge on the same biological output, promoting the increase in cell viability. However, in our experimental system, chemoresistance to CPT-11 appears to be mediated primarily by the activation of the Met-dependent signaling pathway.

Regarding the translation of these results to the clinical setting, the *in silico* analysis through online platforms that combine data with information from the stroma of CRC and healthy patients showed that one of the genes overexpressed in tumor stroma and that has a central role in the functional network is TGFβI. Moreover, by functional enrichment, we observed that DEGs between tumor and healthy stroma samples were associated with TGF-β activity. Recently, research on TGFβI has gained great relevance in inflammatory processes and in the development of different types of cancer, including CRC [53,95,96]. It is known that increased synthesis and release of this factor is associated with a poor prognosis in CRC and TGF-β participates in regulating its expression [53,97]. Consistently, in an *in vitro* and *in vivo* study it was demonstrated that the pro-tumor program orchestrated by TGF-β in CRC cells is mediated by TGFβI [98]. Furthermore, another study showed that one of the functions of TGFBI is the stimulation of tumor angiogenesis [98]. Previously we found that the medium derived from CRC cells promotes the formation of HMEC-1 cell tubes; considering that TGF-β1 is present in the TCM derived from these endothelial cells, a positive feedback mechanism between both cell types in response to PTHrP could be possible. To confirm this hypothesis, it would be essential to further study TGFBI in our system.

It is noteworthy that during the functional enrichment analysis, it was observed that the genes which were overexpressed in the CRC stroma, were also linked with the Met signaling pathway. These findings, together with our previous research indicating PTHrP and Met genes overexpression in CRC samples [99], further support the conclusions presented in this work.

Likewise, when the correlation of the gene expression of the markers studied here was evaluated in CRC samples, a positive correlation was found, not only between PTHrP and surface receptors such as PTHR1 and TGFBR1 but also among the factors known to be present in the TME, such as HGF, TGF-β1 and SPARC. With respect to SPARC, the intensity of the correlation between this protein and HGF or TGF-β1 was the highest of all analyzed. The interrelationship of SPARC with PTHrP was previously determined in our laboratory, indicating a strong correlation between both factors in CRC samples and a synergistic effect in the modulatory effect of migration and the expression of EMT-markers on intestinal tumor cells [13]. Since SPARC is involved in the remodeling of the extracellular matrix, it is possible that it collaborates in the extra-cytoplasmic activation of TGF-β in the TME. The positive correlation between SPARC and TGF-β1 was also reported in CRC by Wang and colleagues [44]. Furthermore, it is known that both factors are involved in events associated with an invasive and migratory phenotype in CRC cells, such as EMT [100]. In the case of Met ligand gene expression, although no studies to date have addressed its interrelationship with SPARC in CRC, both HGF and SPARC are macromolecules widely associated with metastatic processes [101,102], so it is highly likely that their expression and activity are related.

The qualitative analysis of PTHR1 and TGFBR1 protein expression was carried out in samples derived from patients with CRC in order to support the results obtained on *in vitro* and *in silico* investigations. Although only a small number of samples were collected, the observations made so far are consistent with findings previously obtained in this work and in our group’s previous publications. The assessment of Kaplan-Meier curves from patients with CRC and simultaneous overexpression of PTHR1, TGFBR1, or both genes showed a diminishing trend towards the overall survival and a significant difference in progression-free time between those with the PTHR1/TGFBR1 genetic signature overexpressed in the primary tumor and those without. Together, these observations suggest that activation of the PTHR1/TGFBR1 signaling axis is linked to poorer clinical outcomes in CRC patients. While individual gene overexpression does not significantly affect overall survival, the combined signatures are associated with reduced disease-free survival and increased risk of recurrence, supporting a cooperative role of these pathways in tumor progression.

PTHrP, TGF-β1, and their respective receptors fulfill a physiological function in normal tissue. Subversion of their synthesis/mechanism of action would act together with other factors to promote changes associated with tumorigenesis, favoring the involvement of other molecules and signaling pathways in later stages of tumor development.

## 4. Materials and Methods

### Reagents and antibodies

Human PTHrP (1-34), Trypan Blue dye were obtained from Sigma-Aldrich Chemical Co (St. Louis, Missouri, USA). Antibodies were purchased from the following sources: anti-Met, anti-phospho Met (Tyr 1234/1235) and anti-CD44 were provided by Cell Signaling Technology (Beverly, Massachusetts, USA); anti-GAPDH, anti-TGFβ1, and neutral red were from Santa Cruz Biotechnology (Santa Cruz, California, USA); anti-TGFβ1R was from Thermo Fisher Scientific (Waltham, USA). SU11274, fibroblast growth factor (FGF) and epidermal growth factor (EGF) were from Sigma-Aldrich Chemical Co. (St. Louis, Missouri, USA). Protein size markers were from Amersham Biosciences (Piscataway, New Jersey, USA), PVDF (polyvinylidene difluoride) membranes and electrochemiluminescence (ECL) detection kit were from Amersham (Little Chalfont, Buckinghamshire, England). The chemotherapeutic drug Irinotecán (CPT-11) was gently provided by Dr. Ariel Zwenger. The stem cell medium (SCM) was acquired in Life Technologies (California, USA).

### Cell lines and culture

HCT116 human colon cancer cells and HMEC-1 human endothelial cells (American Type Culture Collection, Manassas, VA, USA) were grown at 37 ° C, under 5.5% CO2 atmosphere in the air, in Dulbecco’s Modified Eagle Culture Medium (DMEM) containing 10% heat-inactivated and irradiated fetal bovine serum (FBS), 1% non-essential amino acids, penicillin (100 IU/ml), streptomycin (100 mg/ml) and gentamicin (50 mg/ml). The cells were cultured until reaching 80% confluence and then they were deprived of FBS for 2 hours before treatment [14,15].

### Obtention of endothelial-conditioned media

Endothelial-conditioned media were obtained from cultures of HMEC-1 cells that were treated for 16 hours with 10^-8^ M PTHrP (1-34) (TCM, treated conditioned media) or with the cytokine vehicle as a control (CCM, control conditioned media), always maintaining the same cells/volume ratio. The time of exposure was determined according to previous results [13,15]. The CM were collected and centrifuged for 10 minutes at 10.000 rpm to remove cell debris, and the supernatants were stored at −80°C until used for the treatment of HCT116 cells. Each experiment was carried out with an independently obtained CM. To normalize the results, the protein content of the cells, whence the CM were obtained, was quantified by the Bradford colorimetric method [103]. To do this, aliquots of the cell lysate were taken in duplicate; 2.5 ml of Bradford reagent (Coomassie Brilliant Blue G-250 100 mg/L; 4.75% ethanol and 8.5% phosphoric acid) was added, incubated for 5 minutes and absorbances were measured at 595 nm using a Beckman DU530 spectrophotometer. As a standard of known concentration, 1 mg/ml of bovine serum albumin (BSA) was used.

### HCT116 cells treatment

Treatments were performed by incubating cells with TCM or CCM with the addition of 1% FBS, for different times. Where indicated, cells were pre-incubated for 30 min with SU11274 (0.5 μM), a Met inhibitor. In previous works, we confirmed the effectiveness of the dose of SU11274 selected in accordance with the literature [41]. Control conditions were performed by the addition of an equivalent volume of dimethyl sulfoxide (DMSO), the vehicle of the inhibitors.

### Western Blot Analysis

Cells were washed with phosphate-buffered saline (PBS) with 25mM NaF and 1mM Na_3_VO_4_, and lysed in buffer containing 50mM Tris-HCl (pH 7.4), 150mM NaCl, 3mM KCl, 1mM ethylenediaminetetraacetic acid (EDTA), 1% Tween -20.1% Nonidet P-40, 20 µg/ml aprotinin, 20 µg/ml leupeptin, 1 mM phenylmethylsulfonyl fluoride (PMSF), 25 mM NaF and 1 mM Na_3_VO_4_. Lysates were vortexed for 45 seconds, incubated on ice for 10 minutes and then centrifuged at 14000 g and 4 ° C for 15 minutes. The supernatant was collected and protein quantification was performed by the Bradford method [103]. The proteins present in the samples were separated (30 µg/lane) using sodium dodecyl sulfate (SDS)-polyacrylamide gels (8-10% acrylamide) and electrotransferred to hydrophilic PVDF membranes. The membranes were blocked with 5% skim milk in tris buffered saline - Tween (TBS-T) buffer (50mM Tris, pH 7.2-7.4, 200mM NaCl, 0.1% Tween-20), and after incubated over-night with the appropriate dilution of primary antibody (Met #8198 1:3000, p-Met (Tyr1234/1235) #3077 1:500, CD44 #3570 1:1000, anti-TGFβ1 sc-130348 1:500) in TBS-T with 2.5% BSA. After washing, the membranes were incubated with the appropriate dilution of horseradish peroxidase conjugated secondary antibody in TBS-T with 2.5% skim milk. Finally, proteins were revealed using a commercial electrochemiluminescence kit, the bands obtained were digitized densitometrically and quantified by the Image J program.

### Stripping and reprobe membranes

To remove primary and secondary antibodies, membranes were incubated in stripping buffer (62.5 mM Tris–HCl pH 6.8, 2% SDS and 50 mM β-mercaptoethanol) at 55 °C for 30 min in agitation, washed for 10 min in TBS-T (1% Tween-20) and then they were blocked, as indicated in 2.3. After this procedure, it is possible to re-test antibodies in the membranes.

### Trypan blue dye exclusion test

After treatment, the cells were washed with PBS and then incubated with trypsin-EDTA to separate them from the culture plates to which they were attached. The cell suspension was diluted in equal volumes with a Trypan blue stain (1:1). The number of cells per field was counted in a Neubauer chamber; cells excluding dye were considered viable cells. The number of viable cells was calculated according to the following formula:

Total number of viable cells = mean number of viable cells × dilution factor × 104 [104].

### Neutral Red Uptake assay (NRU)

After treatment, cells were washed with pre-warmed PBS and incubated for 3 hours at 37 °C in a humidified 5% CO_2_ atmosphere with a Neutral Red solution prepared according to: 79ml DMEM without FBS + 1ml Neutral Red. After incubation, the medium was removed and the cells were washed again with PBS. Subsequently, the dye retained in the cells was solubilized with a solution of 1% glacial acetic acid, 50% ethanol 96, 49% H_2_O and the absorbance of the solution, proportional to the number of viable cells, was measured at 540 nm. This assay was performed according to previously published protocols [105].

### Measurements of chemotherapeutic drug effects

CPT-11 cytotoxicity was assessed by the Trypan Blue dye exclusion test. HCT116 cells were seeded in a 24-well plate until 80% confluence and then treated with TCM or CCM and/or CPT-11 (10 uM) for triplicates for 24 hours. The dose of this drug was chosen according to previous works [14]. Where indicated, HCT116 were pre-incubated with SU11274 (0.5uM) [41] or DMSO, the vehicle of the inhibitor. After treatment, cells were washed with a PBS buffer and then incubated with Trypsin-EDTA to separate them from the culture substrate. The total number of viable cells was evaluated by Trypan blue and NRU assay, according to the protocols described above.

### Colonosphere formation assay

According to previously described protocols [68-70], this assay was employed to enrich the culture in cells with CSC properties from tumor cell lines. Initially, cells were seeded at 10 cells/well in 96-well ultra-low adhesion plates on SCM, a DMEM/F12 basal medium supplemented with 10 ng/ml of FGF, EGF, 100 U/ml of penicillin and 100 mg/ml of streptomycin. Media were re-supplemented every 3 to 4 days and maintained for 12 days of growth. The colonospheres were visualized under an inverted microscope and each of them was photographed according to the programmed times. The images were then analyzed using the Image J program. The morphological parameters quantified were: major axis, minor axis and aspect ratio (AR) [106,107].

### Protein neutralization

To neutralize TGF-β1, a mouse IgG1 κ monoclonal antibody (sc-130348 3C11, Santa Cruz Biotechnology) was used at a concentration of 5 μg/ml. Mouse IgG1κ was used as an isotype control. The treatment medium was pre-incubated for 20 minutes with the corresponding antibodies. The cells were subsequently treated with this neutralized medium according to protocols previously described [15,108].

### Patients and Clinical specimens

The following research protocol (File number 8610-017183/2018, registry number N° 11.00.18) has the endorsements of the Advisory Commission on Biomedical Research in Human (CAIBSH) and the approval of the Health Ministry of Neuquén, Argentina (provision 0088-18-1-19) and was authorized by the Ethical Committee of the Hospital Interzonal de Graves Dr. José Penna, Bahía Blanca, Buenos Aires, Argentina (File number 8610-017183/2018, Registry of Investigations in Health (RIS) number N° 11.00.18, re-approval date: August 15, 2023).

Specimens were obtained from patients with colorectal adenocarcinoma which were received at the Hospital Interzonal de Graves Dr. José Penna and at the Hospital Provincial de Neuquén from January 1990 to December 2007. Healthy colorectal tissue samples were assigned to the control group. The medical history from each patient was analyzed including information about: sex; age; primary tumor location; histological grade; initial stage (according to TNM staging system provided for American Joint Committee on Cancer - AJCC - and International Union for Cancer Control - UICC). Exclusion criteria: patients under 18 years of age due to the low prevalence of this cancer in the child and youth population [109]; pregnant patients since high levels of PTHrP have been observed in the placenta and in fetal tissues [110]; lactating patients because high levels of PTHrP have been detected in breast tissue and in breast milk and correlates with serum PTHrP levels in these women [111]; patients who have previously developed other types of tumors. All are exclusion criteria at the time of diagnosis. Confidentiality criteria: based on the regulatory provisions referring to confidentiality in clinical investigations, including Law 11,044 and Law 25,326 on data protection, from Argentina; the Guide for Health Research of the National Ministry of Health, the UNESCO Declaration on the Protection of Genetic Data and the Declaration of Helsinki of the World Medical Association (version 2024) and the Council for International Organizations of Medical Sciences (CIOMS).

Paraffin-embedded archival blocks from primary tumors were reviewed by the Hematoxylin-Eosin staining technique; subsequently, representative histological samples were selected of each patient for immunoassays.

### Immunohistochemistry (IHC)

From each paraffin block, tumor sections were sliced and prepared for immunohistochemical detection using the primary monoclonal antibody anti-TGFβR1 (#PA1-38737 1:100), and anti-PTHR1 antibody (sc-12722 1:50) according to the manufacturer’s protocol. The sections were deparaffinized with xylol for 15 minutes and subsequently rehydrated, exposing themselves to decreasing alcoholic concentrations. Antigenic recovery was carried out using a pressure cooker with sodium citrate buffer (10 mM, pH 6) for 15 minutes at 1 atm. The sections were washed 2 times with PBS and we proceeded to block endogenous peroxidase with H2O2 30% for 10 minutes. After two washes with PBS, the corresponding antibody was added to each sample and incubated over-night at 4 ° C. Immunohistochemical staining was carried out manually using ABCAM Detection IHC Kit (Abcam, Cambridge, MA, USA) according to the manufacturer’s instructions. The images obtained by optical microscope were analyzed using an open source image processing package based on Image J [112,113].

### Microarray gene expression data

Gene expression data from normal samples, adenomas, and colorectal tumors were obtained from the Gene Expression Omnibus (NCBI-GEO) platform. The GSE31279 accession (accessed on 10 February 2024), was queried based on the Illumina humanRef-8 v2.0 platform. Samples were obtained from colorectal cancer patients from the tumor bank of the University Hospital of Mannheim, Germany. The project was sanctioned by the Ethics Committee of the Faculty of Medicine of the University Hospital of Mannheim and informed consent was obtained from the patients or their spouses when the former died. Of the total samples in the study, for this work 10 samples of tumor stroma and 10 samples of healthy stroma from patients who had undergone surgery between 2002 and 2008 were analyzed. This data can be found here: https://www.ncbi.nlm.nih.gov/geo/query/acc.cgi?acc=GSE31279 [114].

### Obtaining and identifying differentially expressed genes (DEGs)

Differential expression analysis was performed with GEO2R (limma web) using the fold change (FC) criterion. To obtain the differentially expressed genes (DEGs), a threshold of FC>2 (an expression level greater than twofold) corresponding to a Log_2_FC>1 was selected for the overexpressed genes. For downregulated genes, a threshold of FC<1/2 (less than half an expression level) was selected, corresponding to a Log_2_FC<-1. Statistical inference was considered by moderated t-student with a probability of p<0.05 and a probability adjusted by Benjamini and Hochberg of padj<0.05 or false discovery rate (FDR).

In addition, the GEPIA2 (“Gene Expression Profiling Interactive Analysis 2”) database was explored to analyze the clinical implication and the interaction between the genes evaluated in this work.

### Statistical Analyses

Data are shown as mean ± SD from *n* independent experiments (*n* indicated in each figure legend). To determine significant differences between two groups of data, Student’s t-test (two-tailed, equal variance) was carried out. p*<0.05, p**<0.01 and p***<0.001 were considered as statistically significant.

## 5. Conclusions

Based on the results presented in this work and as can be seen in **Figure 15**, we suggest a mechanism by which PTHrP released by neoplastic cells and/or stromal cells, present in the TME, acts at least on the endothelial cells favoring the synthesis and secretion of TGF-β1. Probably, this factor and others present in the CM trigger the aberrant activation of several pathways, including Met signaling, to promote the acquisition of an aggressive phenotype in CRC cells. However, more studies are necessary to: 1) evaluate the presence of HGF or PTHrP in the CM; 2) investigate whether TGF-β1 promotes PTHrP expression on intestinal tumor cells or stromal cells, favoring positive feedback in the TME; 3) evaluate the functional interrelationship of TGF-β1 with other factors present in the CM.

**Figure 15.**
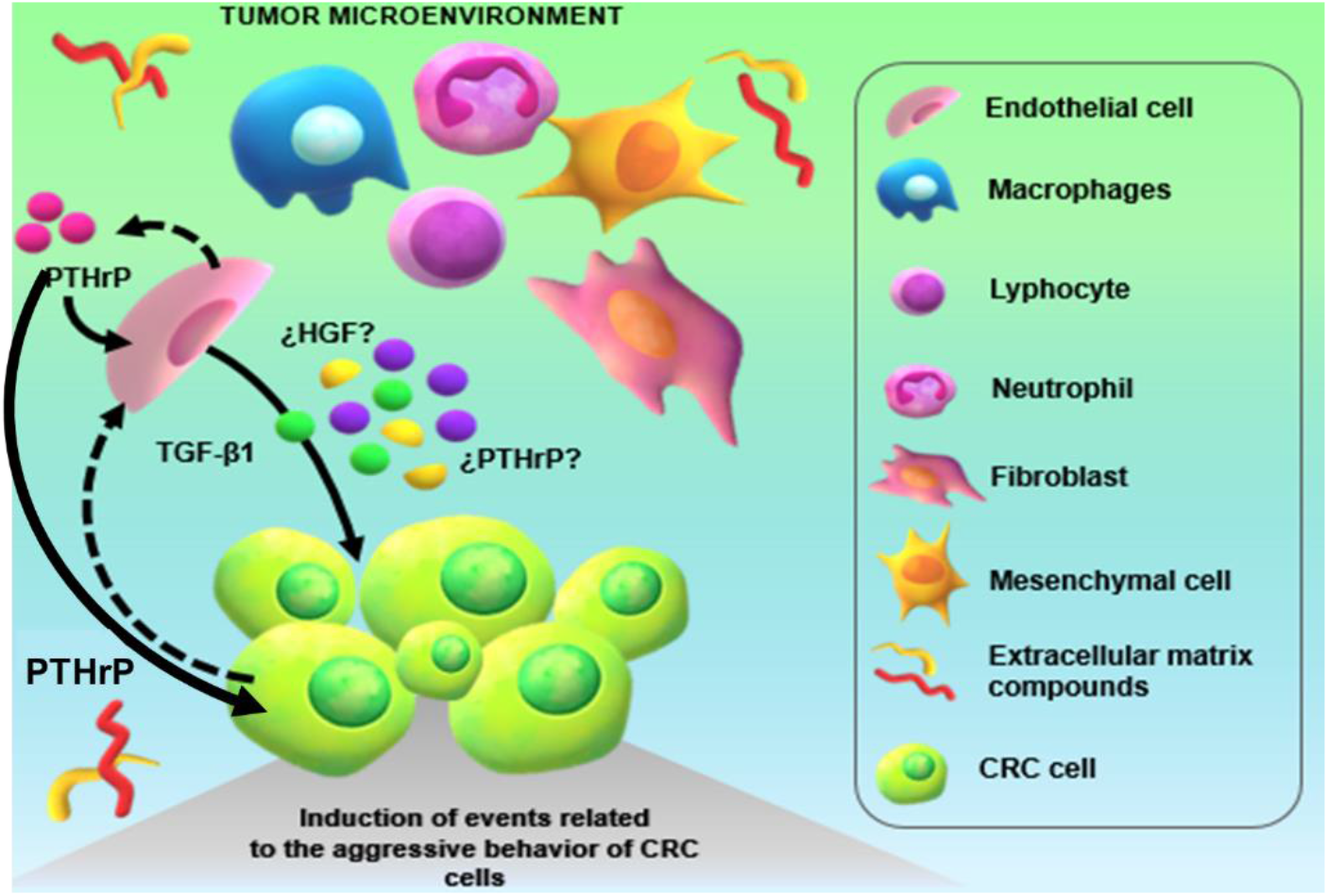
Possible mechanism based on the action of PTHrP on stromal endothelial cells of the TME. PTHrP present in TME, released by neoplastic cells and/or stromal cells, acts at least on endothelial cells, promoting the synthesis and secretion of TGF-β1. Probably this cytokine and other factors present in the TME trigger the aberrant activation of various signaling pathways in CRC cells, favoring the acquisition of an aggressive phenotype, including that of the RTK Met.

## Author Contributions

Conceptualization, C.G. and B.N.; data curation B.N., P.C. and C.G.; formal analysis, B.N., P.C. and C.G; funding acquisition, C.G. and H.C.; investigation, B.N., P.C and C.G.; methodology, B.N., P.C. and C.G.; project administration, C.G.; resources, C.G.; software, P.C.; supervision, C.G. and H.C.; validation, A.Z. and C.G; visualization, P.C., B.N. and C.G; writing-original draft, B.N., P.C., C.B. and C.G.; writing-review and editing, C.G. and H.C. All authors have read and agreed to the published version of the manuscript.

## Institutional Review Board Statement

The study was conducted in accordance with the Declaration of Helsinki, and approved by the Advisory Commission on Biomedical Research in Beings Human Rights (CAIBSH) and the approval of the Health Ministry of the Province of Neuquén, Argentina (provision 0088-18-1-19) and was authorized by the Ethical Committee of the Hospital Interzonal de Graves y Agudos Dr. José Penna, Buenos Aires, Argentina (File number 8610-017183/2018, Registry of Investigations in Health (RIS) number N° 11.00.18, re-approval date: August 15, 2023).

## Informed Consent Statement

Informed consent is waived for each patient due to the sample collection period (1999 to 2007). The non-use of informed consent is justified by article 10 point 3, subsection of the provincial law 15.462 (Buenos Aires, Agentina) in accordance with Council for International Organizations of Medical Sciences (CIOMS) and WMA Declaration of Helsinki – Ethical Principles for Medical Research Involving Human Subjects (2024).

## Data Availability Statement

Datasets are available on request. The raw data supporting the conclusions of this article will be made available by the authors, without undue reservation.

## Acknowledgments

The authors thank Diego Nabaes Jodar for advice on the statistical analysis of the manuscript.

## Disclaimer/Publisher’s Note

The statements, opinions and data contained in all publications are solely those of the individual author(s) and contributor(s) and not of MDPI and/or the editor(s). MDPI and/or the editor(s) disclaim responsibility for any injury to people or property resulting from any ideas, methods, instructions or products referred to in the content.

