## Supplementary material for "PTHrP drives aggressive traits in colorectal cancer cells: Implications of tumor-stromal cells"

*Article*

**Supplementary Material**

**Table 1.** Differentially expressed genes between CRC stromal tissue vs. healthy stromal tissue.

| **Regulation** | **DEG** |
| --- | --- |
| Positive | RAB20; CPZ; PRDX4; RPN1; BICD1; TBX2; PGD; TTYH3; SIM2; APMAP; PLEKHB2; ICA1  SPHK1; TSKU; ITPR3; FAM156A; FCER1G; WBSCR22; OAS1; PLXNA3; PPIB; GALNT5; ELF4; PRKCD; FOXP1; WDR5; APEX2; CTTN  BMP6; MFGE8; SDF2L1; PNKD; CAPG; SOX4  SDSL; BCL2L1; PYCR1; SPINT1; CARHSP1; TMEM200A; CXorf38; MARCKSL1; GLYCTK  WWC1; ID3; CLRN3; LRRC15; CXCL8; SALL4; SMOX; TNFRSF10B; CCNO; PDIA4; TMSB10  AGT; SYK; RNF43; PDGFRB; TMEM2; LRRC32; HLA-E; TMEM74B; NUAK1; DOK4; TNPO1; GNA15; HK3; PLAUR; SPOCD1; EPHA2; APLNR; KDELR3; TESC; MFAP2; EDNRA; C2; **TGFBI;** SH3RF2; LFNG; MXRA5; SLCO4A1; CARD10; HILPDA; ANKRD22; PDPN; TEAD4; SLC1A5; SRPX2; SSR4; SMCO4; CBX2COL1A2; CDC25B; TMEM63A; COL5A2; VASN; GDF15  LAMB3; HOOK1; GJB2; RUNX1; MPZL2; CTSK  SPINT2; PLEK2; KCNJ2; ENC1; ITGA11; CPXM1; COL7A1; DDIT4; BAIAP2L1; FAM150A; CEMIP; ETV4; IER3; MMP1; MARVELD3; C1QTNF5; KRT18; PHLDA2; TRPV4; SFRP2; COL1A1; KCNN4; VCAN; HKDC1; CLDN1; MMP7; CCL20; PLEKHS1; NOX4; DCBLD1; IL1RN; MAPK13; MYEOV; PLAU; S100P; CTHRC1; DIO2; COL8A1; COL11A1; CEACAM6 |
| Negative | PRKCB; SMARCE1; UBE2L3; NMT2; HIRIP3; MIR4697HG; C1orf21; NKIRAS1; SCARA3; DSTN; PMP2; PYGM; KLF9; SGCE; ZSCAN31  DDX46; GFRA2; COX20; GLIPR2; NAP1L5; RIMKLB; FBXL5; ACACB; SH3BGRL; SCARA3; S100B; MOB4; NLGN1; EPM2A; MIR600HG; TBC1D1; PRKCA; AGTR1; LARGE1; FECH; ZDHHC11; GNAZ; ENAH; RSU1; JAM3; DEPTOR; PRICKLE2; MPP2; PELI2; PLA2G5; IFT57; CD36; RAVER2; SORCS1; SSBP2  CDC42EP4; NME4; RERG; KLHL42; DNAJC27; MAP6; ARHGAP10; CELF2; GFRA3; MAOB; NME5; AP1S2; ARNT2; FNBP1; PPP1CB; CAB39L; TSHZ1; KCNMA1; FZD7; GPER1; BCHE; TOX; LIMS2; LRRC49; SYNGR1; OLFML2A; ATP2A2; MEIS2; BCL2L2; C3orf70; CCDC69; ADARB1; CAMK2G; FAM107A; KIZ; FAM172A; GNG3; ARHGEF37; NTNG1; MTURN; LPAR1; PIPOX, CRTAP; KCNIP3; CHODL; MAL; NRXN2; RAB6B; MATN2; GATS; IGSF11; MYL9; SCRG1; SVIL; BVES; SOX10  SPTBN1; SPARCL1; PED1; GPM6B; RIMS3; GHR; DLG2; SLC25A23; CIDEA; SRPX; SMTN; FAXDC2; PDE5A; FAM129A; SPEG; MYOT; RFX2; DTNA; CXCL12; INA; SORBS1; LMOD1; FOXF2; SLC8A2; SOX15; SDPR; HMP19; RALYL; FHL1; SLC6A16; ABCA8; SNAP91; ADAMTSL3; CAP2; ANGPTL1; TMOD1; SORBS2; PDZRN4; AFF3; DPP6; RBPMS2; FRAS1; ABI3BP; ADH1B; PRUNE2; FAM46B; CTXN1; PLCD4; GPM6B; GSTM3; PDK4; PGM5; NNAT; HSPB8; MAMDC2; NPTX1; C7; LMO3; C2orf40; HSPB6; CASQ2; PTGS1; ATP1A2; REEP1; TCEAL2; PNCK |
